# Comparative genomics of pesticide-degrading *Enterococcus* symbionts of *Spodoptera frugiperda* (Lepidoptera: Noctuidae) leads to the identification of two new species and the reappraisal of insect-associated *Enterococcus* species

**DOI:** 10.1101/2023.04.20.537713

**Authors:** Ana Flávia Freitas Gomes, Luís Gustavo de Almeida, Fernando Luis Cônsoli

## Abstract

*Enterococcus* species have been described as core members of the microbial community of *Spodoptera frugiperda* (Lepidoptera:Noctuidae) and have been reported in previous studies as insecticide degrading agents. Phenotypic assays and comparative genomics analyses of several pesticide-degrading *Enterococcus* isolated from the larval gut of *S. frugiperda* led to the identification of *Enterococcus entomosocium* n. sp. and *Enterococcus spodopteracolus* n. sp. Their identities as new species were confirmed by whole genome alignment using the cut-offs of 95-96% for the average nucleotide identity (ANI) and 70% for the digital DNA:DNA hybridization (dDDH) values. The systematic positioning of these new species within the genus *Enterococcus* was resolved using genome-based analysis, placing *Enterococcus casseliflavus* as the sister group of *E. entomosocium* n. sp., and *Enterococcus mundtii* of *E. spodopteracolus* n. sp. Comparative genomic analyses of several isolates of *E. entomosocium* n. sp. and *E. spodopteracolus* n. sp. led to a better assessment of the interactions established in the symbiotic association with *S. frugiperda*, and the discovery of misidentified new species of *Enterococcus* associated with insects. Our analyses also indicated the potential of *E. entomosocium* n. sp. And *E. spodopteracolus* n. sp. to metabolize different pesticides arises from molecular mechanisms that result in the rapid evolution of new phenotypes in response to environmental stressors; in this case, the pesticides their host insect is exposed to.

## Introduction

Current research into the functional contribution of host-associated bacteria has demonstrated their participation in host phenotype definition, which includes various types of disfunctions in humans [1, 2] to changes in body coloration in aphid insects [3]. The diversity of associations and contributions of microorganisms to host insects is almost as diverse as insects are. Insect-associated microbes directly interfere with the host response to a range of biotic and abiotic stressors, contributing to host fitness [4–6]. Microbes may alter insect adaptiveness through nutritional regulation, nutrient contribution, digestion of recalcitrant substrates, hormonal signaling, immune contribution, and xenobiotic metabolism [4, 7, 8].

The megadiverse order Lepidoptera has a long coevolutionary history with plants and is almost entirely associated with angiosperms [9]. The divergence of the major ditrysian lineages in the Cretaceous and the explosion of lepidopterans nearly 90 mya [10] resulted in the second largest group of phytophagous insects [9]. Several lepidopterans are of great importance to man, particularly by the risk they pose to food security, acting as severe pests of many staple foods and other agricultural products [11].

Lepidopterans are also known to host a range of microbes, with bacteria mainly inhabiting the gut lumen [12, 13]. Early studies on the microbial association with the lepidopteran gut disbelieved the existence of intimate microbial partnerships [14], but several other studies with lepidopteran larvae have demonstrated that gut bacteria can aid in food digestion [15], provide nutrients [16] and antimicrobials to regulate pathogens in the host gut microbiota [17] and participate in the metabolization of host plant-derived defense molecules [18, 19] and pesticides [20–22].

*Proteobacteria* and *Firmicutes* dominate the gut microbiota of insects, with relative proportions reflecting host-insect groups [23, 24]. *Enterococcaceae* is one of the most common *Firmicutes* in insect gut microbial communities [24, 25], and the genus *Enterococcus* has been reported to occur persistently in several species of lepidopterans regardless of diet and insect metamorphosis (reviewed by [12]). *Enterococcus* have also been reported to contribute to the metabolization of xenobiotics in lepidopterans [26, 27], including *Spodoptera* larvae [28, 29]. Its prevalence in the larval gut microbiota of *Spodoptera littoralis* and *Spodoptera frugiperda* recognized *Enterococcus* as a core member of their microbial community [25, 30, 31]. However, there are associations in which they were reported to establish pathogenic interactions with lepidopteran host larvae [32–34]. In *Spodoptera*, *Enterococcus faecalis* has been shown pathogenic to *Spodoptera exigua* [35], and *Enterococcus mundtii* [36] and *Enterococcus cloacae* [37] to *Spodoptera litura*.

Many *Enterococcus* colonies were isolated from the gut of several laboratory-selected insecticide-resistant and susceptible strains of *S. frugiperda.* The isolates obtained from insecticide-resistant strains of *S. frugiperda* were capable to grow on selective media containing the insecticide by which host insect was selected for resistance, but not the isolates obtained from susceptible larvae [28]. These isolates were putatively identified as *E. mundtii* and *Enterococcus casseliflavus* by heuristic comparisons and phylogenetic analysis based on an almost complete sequence of the 16S rRNA gene [28].

The diversity of interactions that *Enterococcus*, including *E. mundtii* and *E. casseliflavus*, establish with their host insects [38], and the great diversity in the potential for insecticide degradation of the putative *E. mundtii* and *E. casseliflavus* isolates obtained from insecticide-resistant strains of *S. frugiperda,* led us to conduct phenotypic assays and apply comparative genomic analyses to look into the physiological diversity and the type of interaction these *Enterococcus* hold with the larvae of *S. frugiperda,* bringing to light their potential to metabolize insecticides and the relevant features they carry as symbionts.

## Methods

### Bacterial isolates

Insecticide-degrading isolates putatively identified as *E. casseliflavus* or *E. mundtii* obtained from the larval gut of *Spodoptera frugiperda* resistant to insecticides [28] were selected for this study (Table 1). Additional isolates of *E. casseliflavus* and *E. mundtii* that were unable to grow in insecticide-selective media were also isolated from the larval gut of a susceptible reference line of *S. frugiperda* that has been maintained without any source of insecticide selection pressure for over 15 years (Table 1).

**Table 1.**
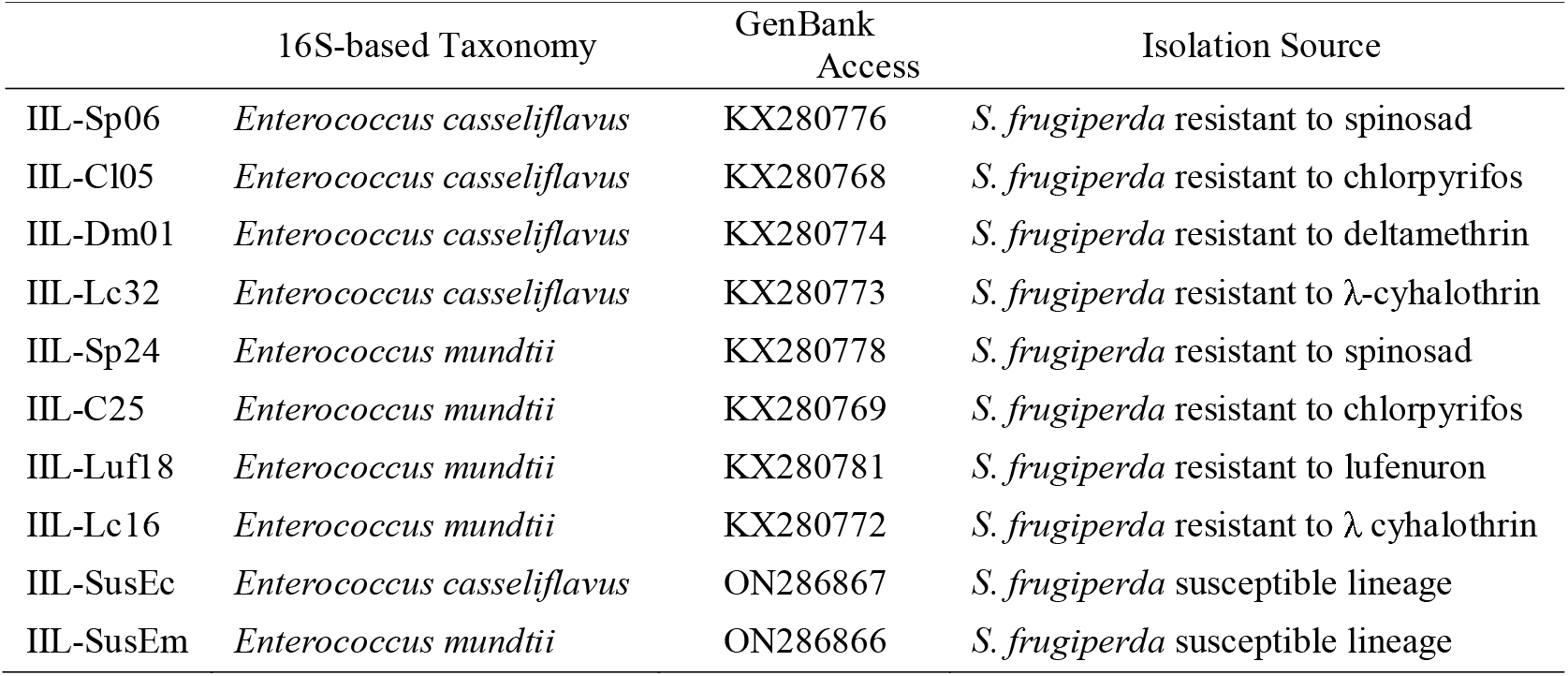
Bacterial phylotypes with differential capacity to metabolize insecticides compounds. Strains isolated from susceptible and insecticide-resistant larvae of Spodoptera frugiperda.

### Isolate phenotyping

#### Antibiotic susceptibility

Antibiotic susceptibility testing was performed using the disk diffusion technique [39] with a commercial system for testing bacterial response to several antibiotics (Antibiotic Multidisc - Laborclin, Brazil). The isolates IIL-Cl05, IIL-Dm01, IIL-Lc32 and IIL-Sp06, putatively identified as *E. casseliflavus*, and the isolates IIL-Cl25, IIL-Lc16, IIL-Luf18 and IILSp24, putatively identified as *E. mundtii* were grown in TSB medium under constant shaking (120 rpm at 28°C) to OD_600_ = 0.450. Cultures were subjected to micro-colony assays and adjusted to an initial cell density of 10^6^ CFU/mL. The isolates were seeded (0.5 mL) in Petri dishes (150 x 15 mm) on TSA medium. Afterwards, the multi-discs ring of antibiotics was set onto the seeded medium. Plates were incubated at 28°C and the susceptibility to the antibiotics assessed by measuring the inhibition zone (in mm) after 48 h of incubation at 28°C. The antibiotics used were cefepime (Cpm-30µg), ciprofloxacin (Cip-5µg), clindamycin (Cli-2µg), chloramphenicol (Clo-30µg), erythromycin (Eri-15µg), gentamicin (Gen-10µg), oxacillin (Oxa-1µg), penicillin (Pen-10U), rifampin (Rif-30µg), sulfazotrim (Sut-25µg), tetracycline (Tet-30µg), and vancomycin (Van-30µg).

#### Carbohydrate metabolism

The selected isolates were also tested for their capacity to metabolize 49 carbohydrates using the API 50 CH system (BioMerieux, France), following the manufacturer’s recommendations. An initial cell suspension at a density of 10_ CFU/mL was prepared as before. Cells were pelleted by centrifugation (1000 *g* x 2 min x 20°C) and resuspended in an appropriate medium composed of ammonium sulfate (2 g), yeast extract (0.5 g), tryptone (1 g), disodium phosphate (3.22 g), potassium dihydrogen phosphate (0.12 g), and phenol red (0.17 g) in demineralized water at pH 7.4 (1000 mL). The cell suspension obtained was equally distributed in the available microtubes for the metabolism trials. Microtubes were incubated at 28°C and evaluated after 48h. Positive results were indicated by changes in the medium color due to the acid production resulting from metabolization of one of the following carbohydrates: glycerol, erythritol, D-arabinose, L-arabinose, D-ribose, D-xylose, L-xylose, D-adonitol, methyl-ß-D-xylopyranoside, D-galactose, D-glucose, D-fructose, D-mannose, L-sorbose, L-rhamnose, dulcitol, inositol, D-mannitol, D-sorbitol, methyl-α-D-mannopyranoside, methyl-α-D-glucopyranoside, N-acetylglucosamine, amygdalin, arbutin, esculin ferric citrate, salicin, D-cellobiose, D-maltose, D-lactose (bovine origin), D-melibiose, D-saccharose (sucrose), D-trehalose, inulin, D-melezitose, D-raffinose, starch, glycogen, xylitol, gentiobiose, D-turanose, D-lyxose, D-tagatose, D-fucose, L-fucose, D-arabitol, L-arabitol, potassium gluconate, potassium 2-ketogluconate, or potassium 5-ketogluconate.

### Comparative genomic analysis

#### DNA extraction and sequencing

Bacterial isolates were cultivated in TSB medium under constant agitation (120 rpm x 28 °C x 24 h). Cells were pelleted (2000 *g* x 5 min) and subjected to genomic DNA (gDNA) extraction using the Wizard Genomic DNA Purification kit (Promega), according to the manufacturer’s recommendations. gDNA quality, integrity and purity was verified by electrophoresis in a 0.8% agarose gel slab containing 0.5 μg/mL of ethidium bromide in TAE buffer (40 mM Tris-acetate, 1 mM EDTA, pH 8.2) under constant voltage (90V), and by determination of UV absorbance ratio (Abs_280_/Abs_260_). The extracted gDNA was sent to the Multiuser Center for Agricultural Biotechnology at USP/Esalq for library preparation and sequencing on an Illumina MiSeq v3 platform using a paired-end strategy (2x300bp).

#### Genome assembly and annotation

Sequence reads were screened using FastQC 0.10.1 [40], and quality filters were applied to eliminate residual Illumina adapters (PE-Nextera), low-quality bases (Phred Quality Score < 30) and reads shorter than 25 bp using Trimmomatic-0.39 [41]. Genomes were assembled based on SPAdes 3.15.4 pipeline for isolate bacterial datasets [42, 43] using both paired and orphan reads. Genomes are available at the National Center for Biotechnology Information (https://www.ncbi.nlm.nih.gov/) under bioproject PRJNA943201(accession numbers SAMN33713714-SAMN33713718) and PRJNA944113 (accession numbers SAMN33735643-SAMN33735646). Plasmids were detected using MegaBLAST annotation (BLAST+ version 2.13.0) and were filtered from the analysis using CLC Genomics Workbench v21.0.5 (QIAGEN).

Draft genomes and filtered plasmids were separately annotated at the Bacterial and Viral Bioinformatics Resource Center (BV-BRC) website (https://www.bv-brc.org/) using the Rapid Annotation Subsystem Technology toolkit (RASTtk) [44]. Features related to assembly quality, identification, and assignment of sequences into functional categories (subsystems) were included in all genome annotations. The parameters coarse consistency (CC), fine consistency (FC), completeness and percentage of contamination, as well as the number and size of contigs were used to estimate the assembly’s quality. Proteins assigned to functional categories were characterized according to the Enzyme Commission (EC) numbers and Gene Ontology (GO), and whenever possible mapped to KEEG pathways. Specialty genes related to antibiotic resistance, virulence factors, drug targets and drug transporters were annotated using CARD [45], DrugBank [46], TCDB [47], VFDB [48] and VICTORS [49] databases as implemented in BV-BRC (https://www.bv-brc.org/).

Biosynthetic gene clusters (BGCs) were predicted with AntiSMASH (https://antismash.secondarymetabolites.org/) [50]. Well-defined clusters containing all required parts and partial clusters missing one or more functional parts (relaxed mode of detection strictness) were searched, and the results were visualized using the output files from AntiSMASH. BGCs type and organization were analyzed, as well as the percentage of gene similarity with the closest known compound from MiBIG database (https://mibig.secondarymetabolites.org/) in the procluster-to-region mode [51].

#### Whole genome alignment, ANI and dDDH analyses

The assembled genomes were aligned against type strains of *E. casseliflavus* EC20^T^ (GenBank CP004856.1) and *E. mundtii* Qu25 (GenBank: AP013036.1) using the genomic tools available in CLC Genomics Workbench (QIAGEN) v21.0.5. Contigs were reordered according to the reference genome used. Genomic alignments were conducted using an initial seed length of 15 bp and a minimum alignment block length of 100 bp. The similarity of the aligned sequences was quantified according to the percentage of genome alignment (AP) and the average nucleotide identity (ANI).

Genome-based taxonomy was inferred through digital DNA: DNA hybridization (dDDH) in the Type (Strain) Genome Server (TYGS) (https://tygs.dsmz.de/) [52]. Pairwise dDDH values between the investigated phylotypes and type-strains were achieved by the sum of all identities found in high-scoring segment pairs (HSP) divided by overall HSP length (TYGS d4 formula). The cut-off of 70% for dDDH and 95∼96% ANI were adopted for species delineation [53, 54]. Subspecies clustering was done using a dDDH cut-off value of 79% [55].

#### Molecular markers for species differentiation

Bacterial phylotyping was carried out by comparing the house-keeping genes ATP synthase alpha chain (atpA), D-alanine-D-alanine ligase (ddl), glutamate dehydrogenase (glucose-6-phosphate 1-dehydrogenase) (gdh), phosphate-binding protein (pstS), N5-carboxyaminoimidazole ribonucleotide synthase (purK), and xanthine phosphoribosyltransferase (xpt) which are used in the characterization of Enterococcus faecalis and Enterococcus faecium as available in MLST database (https://pubmlst.org/databases/). Comparisons were conducted using only the gene region proposed as representative for typing these species.

Sequences of the selected isolates and of reference strains were retrieved based on the RASTtk annotation at BV-BRC website and were analyzed by clustalW alignments (gap opening penalty=15, gap extension penalty=6.66) using MEGA11 [56]. The phylogenetic tree for each group of gene sequence was constructed using the GTR, Tamura 3-parameter or Tamura-Nei substitution models, and the phylogenies were tested using the maximum likelihood method. The selection of the evolutionary models was based on the Akaike and the Bayesian information criteria for the goodness of fit obtained for each set of data. The proportion of replicate phylogenies were estimated by bootstrap analysis with 500 iterations. All phylogenetic analyses were done in MEGA11 [56]. The strains *Enterococcus avium* ATCC 14025 (GenBank ASWL00000000), *Enterococcus durans* KLDS6.0930 (GenBank CP012384.1), *Enterococcus casseliflavus* EC20 (GenBank CP004856.1), *Enterococcus faecalis* V583 (GenBank AE016830.1), *Enterococcus gallinarum* EG2 (GenBank ACAJ00000000.1), *Enterococcus mundtii* Qu25 (GenBank AP013036.1), and *Enterococcus termitis* LMG 8895 (GenBank MIJY00000000.1) were included in the phylogenetic analysis. *Lactobacillus garvieae* ATCC49156^T^ (GenBank AP009332.1) and *Lactobacillus lactis* UC509.9^T^ (GenBank CP003157.1) were used as external groups.

#### Phylogenetic analysis

The systematic position of the phylotypes of *Enterococcus* cultured from the gut of *S. frugiperda* larvae, as well as the genomic variability among representatives of other *Enterococcus* species were predicted using the whole-proteome-based analysis available in the TYGS server. The phylogenetic tree was inferred with FastME 2.1.6.1 [57] from whole proteome based GBDP distances. Other 28 genomes of *Enterococcus* were used in this analysis (Supplementary Table S1). Alignments were also conducted against 22 additional genomes of strains of *E. casseliflavus* and *E. mundtii* obtained from different isolation sources (vertebrates, invertebrates, soil, and plants), and 17 *Enterococcus* species/strains associated with invertebrates or plants for verification of environmental adaptations in *Enterococcus* (Supplementary Table S2). Protein and nucleotide sequences of 100 PGFams (Supplementary Table S3) were used in *MUSCLE* [58] for protein sequence alignment, and in *Codon_align* in BioPython [59] for nucleotide sequence alignment. The phylogenetic analysis was conducted using the LG model [60] using the *CodonTree analysis* tool as available in BV-BRC. *Lactobacillus garvieae* ATCC49156^T^ (GenBank AP009332.1) and *Lactobacillus lactis* UC509.9^T^ (GenBank CP003157.1) were used as external groups. Both TYGS and BV-BRC phylogenetic trees were supported by bootstrap analysis with 100 iterations. The analysis regarding environmental adaptations in *Enterococcus* was complemented by a digital DNA: DNA hybridization and a whole-proteome-based phylogeny performed on TYGS server.

## Results

### Isolates IIL-Cl05, IIL-Dm01, IIL-Lc32, IIL-Sp06 and IIL-SusEc (putative *Enterococcus casseliflavus*)

#### Antibiotic susceptibility and carbohydrate metabolism phenotyping

No phenotypic differences were observed among the pesticide-degrading isolates IIL-Cl05, IIL-Dm01, IIL-Lc32, and IIL-Sp06 regarding their susceptibility to the tested antibiotics or carbohydrate metabolism. All of them were resistant to oxacillin (1 µg) and tetracycline (30 µg), while being most susceptible to erythromycin (15 µg) (Fig. 1A). They were able to metabolize the same 26 out of the 49 carbohydrates available in the API 50 CH test (Fig. 1A and Supplementary Figure S1).

**Fig. 1.**
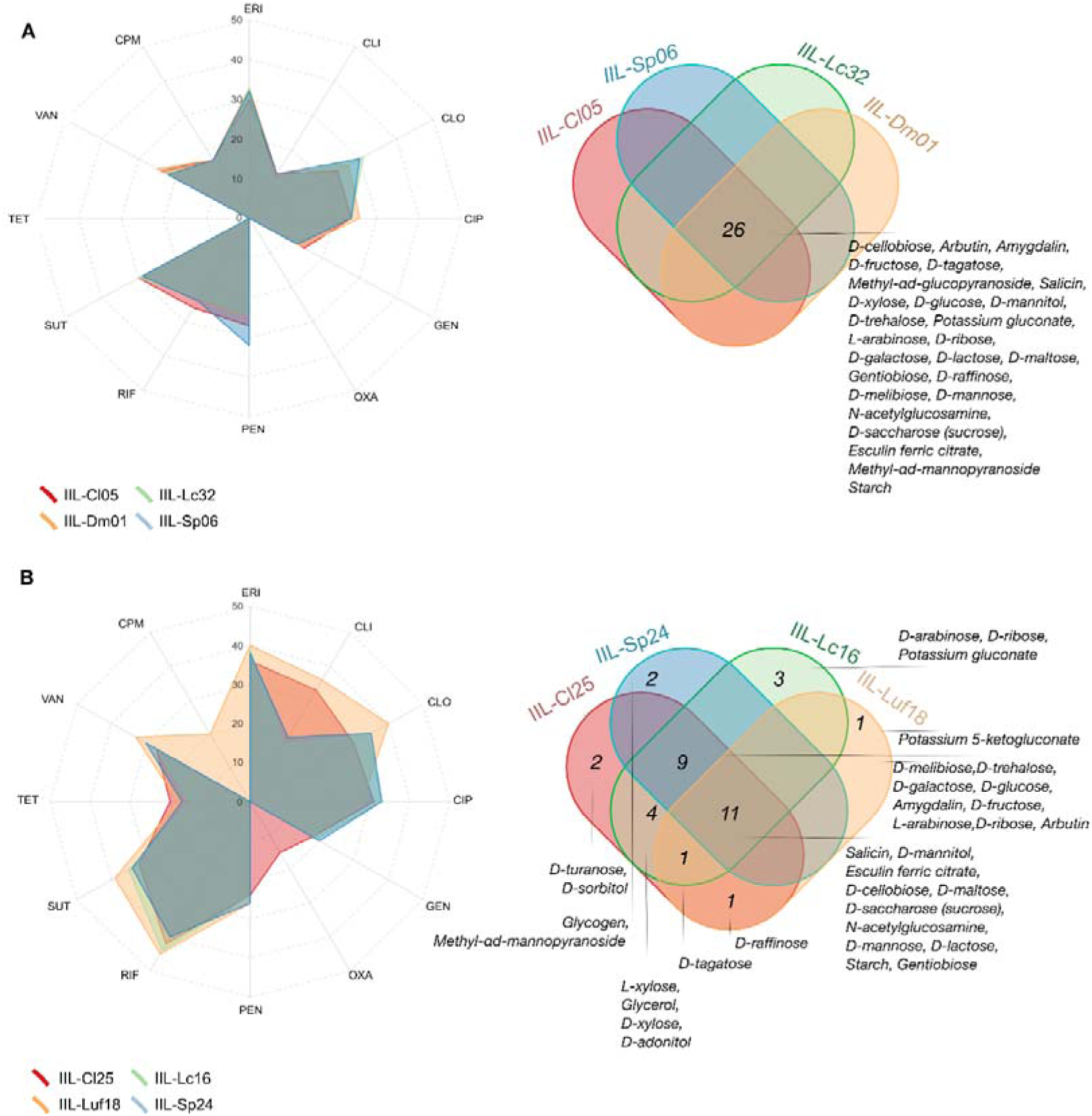
Antibiotic susceptibility (left) and carbohydrate metabolism (right) phenotyping of IIL-Cl05, IIL-Sp06, IIL-Lc32 and IIL-Dm01 (A); and IIL-Cl25, IIL-Sp24, IIL-Lc16 and IIL-Luf18 (B). Antibiotic susceptibility evaluation to the compounds cefepime (CPM-30µg), ciprofloxacin (CIP-5µg), clindamycin (CLI-2µg), chloramphenicol (CLO-30µg), erythromycin (ERI-15µg), gentamicin (GEN-10µg), oxacillin (OXA-1µg), penicillin (PEN-10U), rifampin (RIF-30µg), sulfazotrim (SUT-25µg), tetracycline (TET-30µg), and vancomycin (VAN-30µg). Carbohydrates metabolized in the API 50 CH galleries test.

#### Genome assembly and annotation

The whole genome sequencing of the isolates putatively identified as *E. casseliflavus* from susceptible and insecticide-resistant *S. frugiperda* larvae generated assemblies ranging from 424 to 1173 Mb nucleotides/isolate, with a minimum of 105x and a maximum coverage of 296x. Read trimming and filtering retained around 98% of the reads per library, from which 79-97% were paired reads (Supplementary Table S4).

The resulting assembled draft genomes were in average around 4 Mb in size (42% GC) containing from 131 to 272 contigs, each yielding from 3,818 to 4,087 annotated coding region sequences (CDS). Nearly 66% of all CDS identified were functionally annotated proteins, with 33-34% assigned to EC numbers, 28% to GO terms, and 23-24% mapped to one of KEGG pathways (Fig. 2). Transferases (259) and hydrolases (160), followed by oxidoreductases (97), lyases (69), ligases (64), and isomerases (53) were the major groups of functionally annotated enzymes (Supplementary Table S5). Most CDS assigned to subsystems were related to *bacterial metabolism* (35%), *protein processing* (16%), and *energetic processes* (15%). The remaining CDS were mainly linked to the *stress response, defense,* and *virulence* subsystem or to those subsystems associated with *cellular* and *DNA processes* (Fig. 2). IIL-Sp06 was the isolate with the greatest number of proteins assigned to the subsystem *prophages, transposable elements, and plasmids* (Fig. 2).

**Fig. 2.**
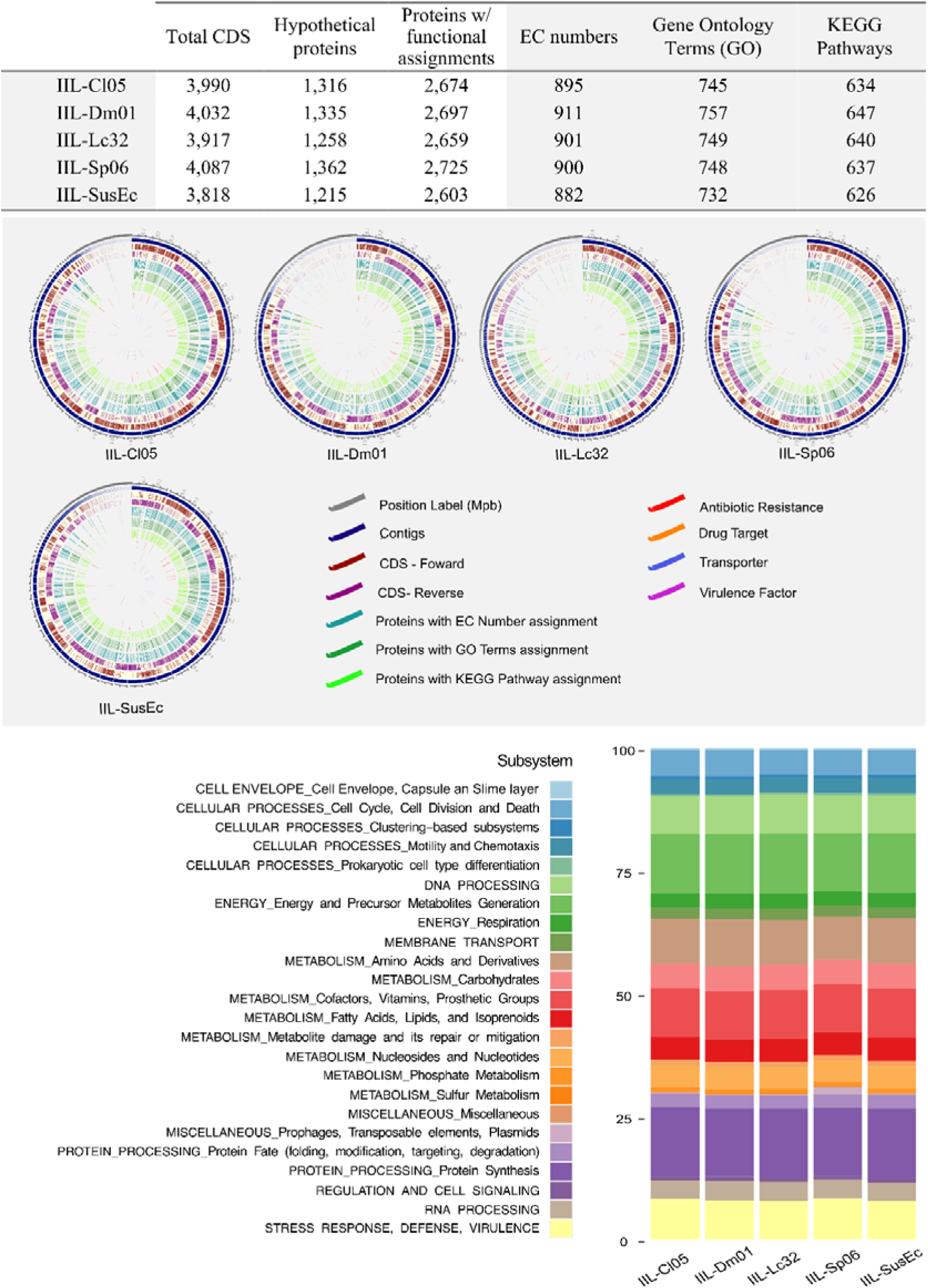
Genomes annotation of IIL-Cl05, IIL-Sp06, IIL-Lc32, IIL-Dm01 and IIL-SusEc by Rapid Annotation Subsystem Technology toolkit at BV-BRC website.

Most sequences mapped to KEEG pathways were functionally related to *carbohydrate metabolism* (29%), *amino acid metabolism* (14%) or *energy metabolism* (11%) (Fig. 3). Around 5% of the mapped sequences were associated with *xenobiotics biodegradation and metabolism*, such as the ones associated with the metabolism of xenobiotics by cytochrome P450, and with the degradation of compounds as gamma-hexachlorocyclohexane, fluorobenzoate, and tetrachloroethene (Fig. 3).

**Fig. 3.**
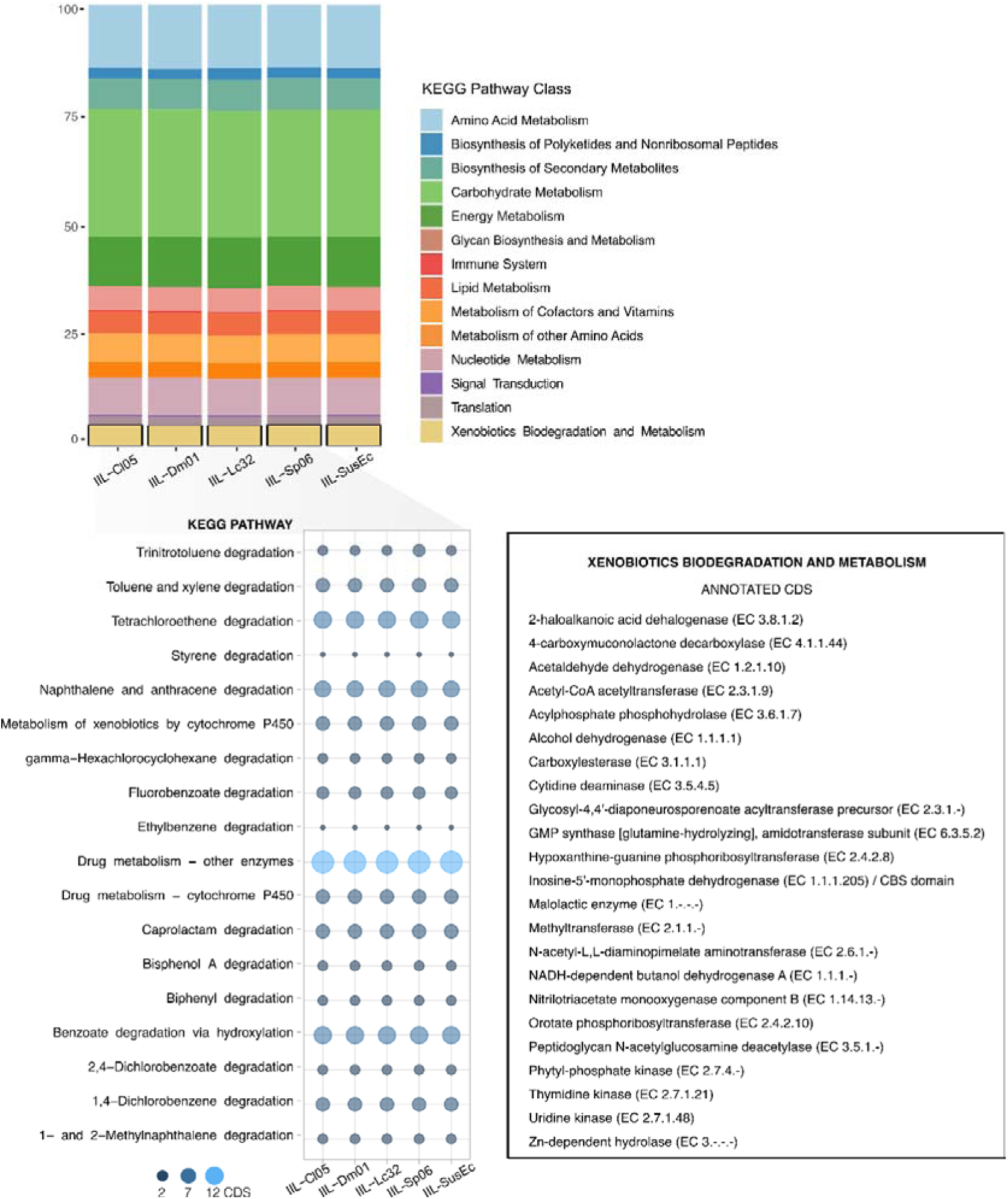
KEGG pathways annotated in the genomes of IIL-Cl05, IIL-Sp06, IIL-Lc32, IIL-Dm01 and IIL-SusEc. Highlighted, genomic sequences mapped with pathways related to xenobiotics biodegradation and metabolism.

IIL-Sp06, IIL-Dm01, IIL-Cl05 and IIL-Lc32 from insecticide-resistant lines of *S. frugiperda* and IIL-SusEc from the susceptible line shared a total of 2,125 coding regions (Fig. 4), with an average length of 500 bp and about 87% of functional annotation. IIL-Sp06 was the isolate with the largest number of unique CDS (38), followed by IIL-Lc32 (18) and IIL-Dm01 (17). IIL-Sp06 shared 23 CDS only with IIL-Cl05, and 14 only with IIL-Dm01. IIL-Dm01 and IIL-Sp06 also shared 11 coding regions with IIL-Cl05. No unique CDS was found in the IIL-SusEc genome.

**Fig. 4.**
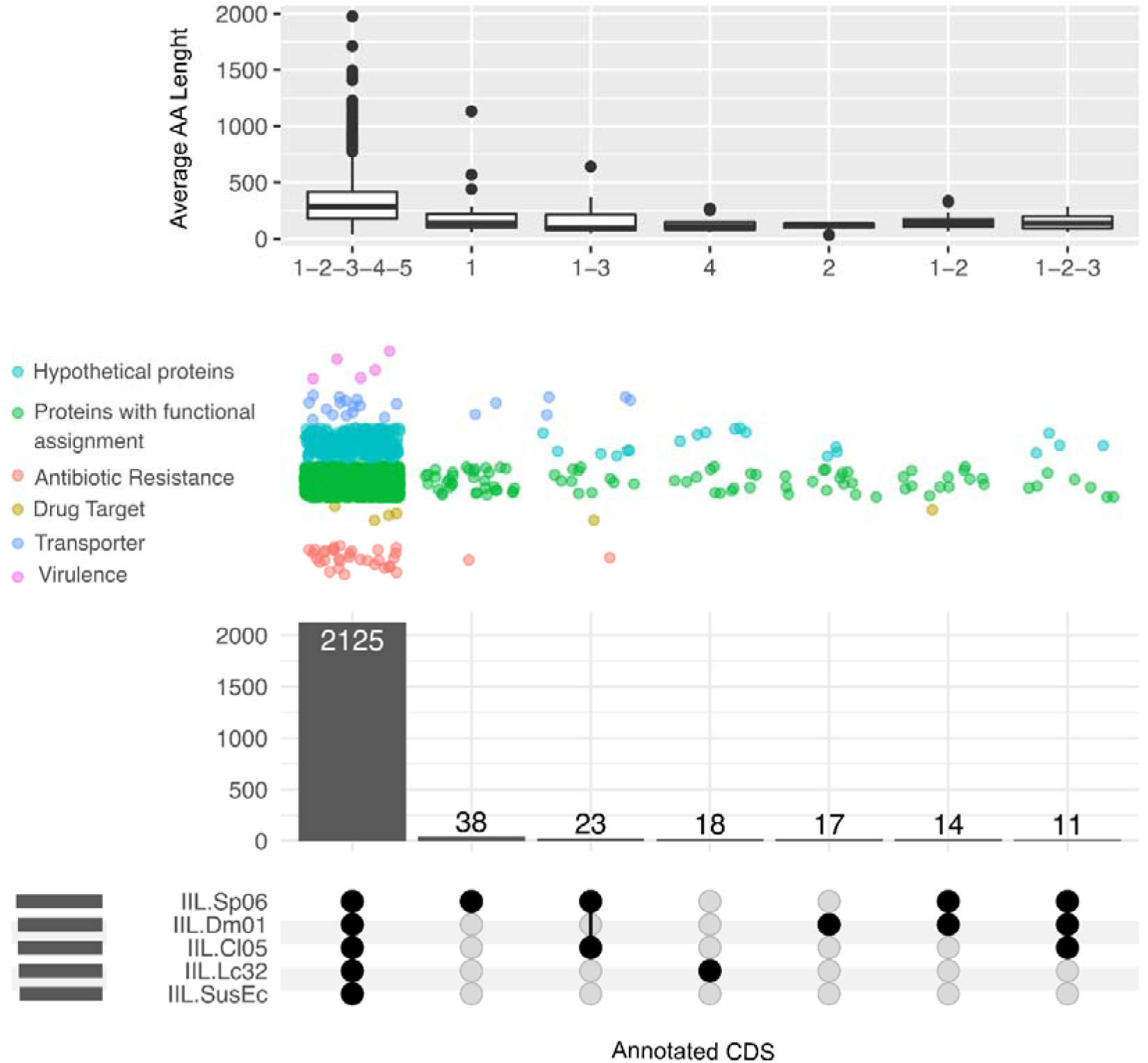
Shared coding regions sequences (CDS) between the isolates IIL-Sp06, IIL-Dm01, IIL-Cl05, IIL-Lc32 and IIL-SusEc. Classification and average amino acid string length of each annotated CDS.

Most of the antibiotic resistance genes (96%) identified were common to all isolates, which included antibiotic inactivation enzymes, antibiotic target in susceptible species, antibiotic target modifying/replacement enzymes, and regulators of antibiotic resistance gene expression, among others (Supplementary Table S6). The tetracycline resistance ribosomal protection protein Tet(S) was annotated only in the genomes of IIL-Sp06 and IIL-Cl05, and a class A broad-spectrum beta-lactamase (EC 3.5.2.6) was exclusively annotated in IIL-Sp06 (Supplementary Table S6).

The drug target regions *DNA gyrase subunit B* (EC 5.6.2.2), *IMP dehydrogenase* (EC 1.1.1.205), *LSU ribosomal protein L3p*, and the *phosphocarrier protein Hpr* were annotated in the genome of all isolates. Two phage-related proteins, however, were detected only in IIL-Cl05, IIL-Dm01, and IIL-Sp06. The *phage recombination protein NinB* was detected in IIL-Sp06 and IIL-Dm01, while the *phage lysozyme R* (EC 3.2.1.17) was present only in IIL-Sp06 and IIL-Cl05.

All isolates shared from 89 to 100% of the proteins annotated as transporters (Supplementary Table S6), most of which classified as a heterodimeric efflux pump from the *atp-binding cassette (abc) superfamily* or as membrane proteins from *nhac Na(+):H (+) antiporter (nhac)* family. The multidrug resistance efflux pump *PmrA*, as well as the multidrug-efflux transporter transcription regulator *BltR* were also annotated in the genome of all isolates. For unique transporters, stands out those from the *type iv (conjugal dna-protein transfer or virb) secretory pathway (ivsp) family*, annotated in IIL-Cl05 and IIL-Sp06. The virulence factors *amidine-lyases* (EC 4.3.2.2), *oxidoreductases* (EC.1.1.1), *methionine aminopeptidases* (EC 3.4.11.18), and the *translation elongation factor LepA* were common to all isolates.

Plasmids sequences filtered from the draft genomes obtained were putatively identified by RASTtk annotation (Supplementary Table S7). Among the plasmid proteins functionally assigned are *transcriptional regulators*, *antibiotic resistance genes*, and *antitoxins*. BGCs prediction by AntiSMASH detected two clusters encoding biosynthesis proteins classified as *Terpene* and *T3PKS types* (Supplementary Figure S2). Both were found in all isolates. Terpene BGC shared 46% of similarity with a known carotenoid biosynthetic gene cluster from *Rhodobacter sphaeroides* (BGC0000647). T3PKS BGC, however, shared only 22% of similarity with the viguiepinol BGC from *Streptomyces sp.* KO-3988 (BGC0000286).

#### Whole genome alignment, ANI values and dDDH analyses

Whole genome alignment of IIL-Cl05, IIL-Dm01, IIL-Sp06, and IIL-SusEc demonstrated genomes with 96-98% of similarity, and pairwise average nucleotide identities (ANI) of 100% (Fig. 5). But whole genome similarity against the reference *E. casseliflavus* EC20 genome was much lower (80%), and the ANI values obtained were below 95%. Pairwise dDDH analysis agreed with the ANI values obtained for pairwise comparisons between the isolates of *S. frugiperda* and the reference *E. casseliflavus* strain (dDDH value < 70%) indicating the isolates investigated do not belong to known species (Fig. 6 and Supplementary Table S8). Therefore, the partial 16S-rDNA sequence did not provide the necessary information for the correct species identification and phylogenetic placement of the studied isolates (see below). Thus, based on the ANI and dDDH values, we propose the *Enterococcus* isolates studied that share 16S-rDNA sequences identical to *E. casseliflavus* in fact belong to a new species, that we propose to name it as *Enterococcus entomosocium* n. sp., electing isolate IIL-Cl05 as the genome type species.

**Fig. 5.**
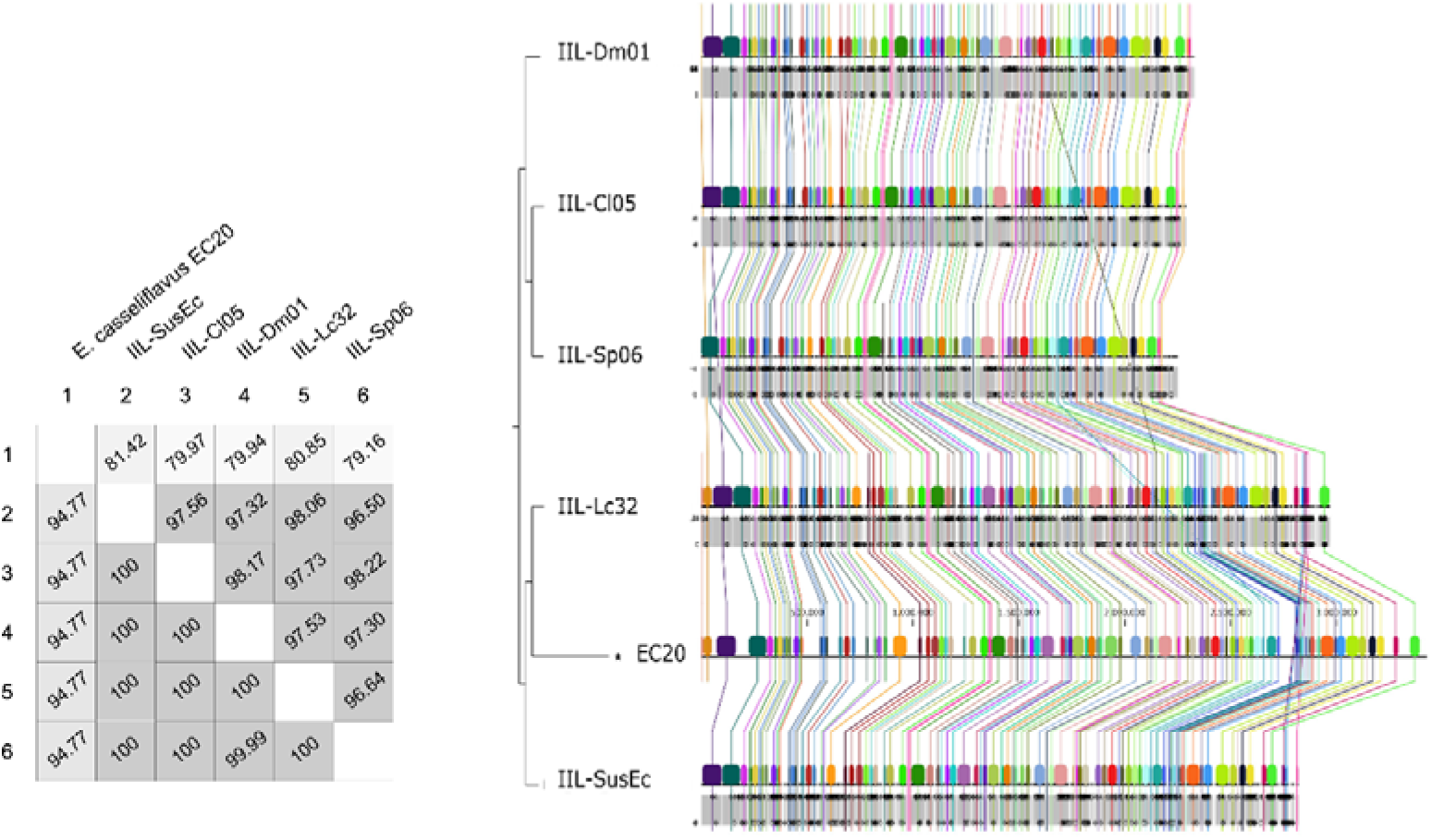
Whole genomes alignment performed at CLC Genomics Workbench (QIAGEN) platform. Percentage of similarity between the isolates IIL-Sp06, IIL-Dm01, IIL-Cl05, IIL-Lc32 and IIL-SusEc, and the reference EC20 (GenBank CP004856.1). In the figure, percentage of alignment (AP) (upper) and the average nucleotide identity (ANI) (down).

**Fig. 6.**
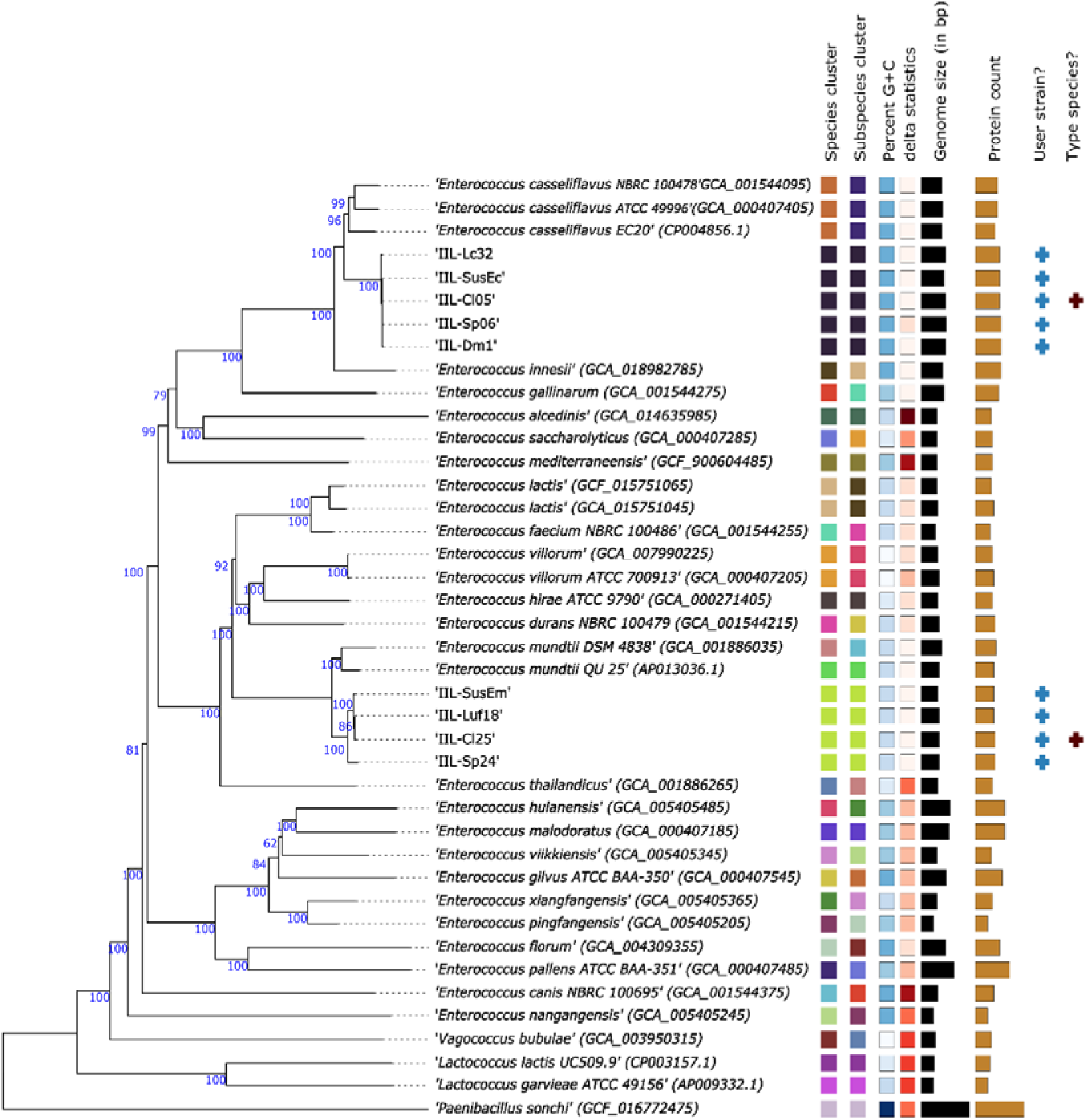
Phylogenetic positioning of isolates of *Enterococcus entomosocium* n. sp. (IIL-Cl05, IIL-Sp06, IIL-Lc32, and IIL-SusEc) and *Enterococcus spodopteracolus* n. sp. (IIL-Cl25, IIL-Sp24, IIL-Luf18, and IIL-SusEm) from resistant and susceptible strains of *Spodoptera frugiperda* within the genus *Enterococcus*. Whole-proteome-based phylogeny performed at the Type (Strain) Genome Server (TYGS). The cut-offs of 70% and 79% for digital DNA: DNA hybridization (dDDH) value were adopted for, respectively, species and subspecies delineation. The values in the different branches correspond to bootstrap values.

### Isolates IIL-Cl25, IIL-Luf18, IIL-Sp24, and IIL-SusEm (putative *Enterococcus mundtii*)

#### Antibiotic susceptibility and carbohydrate metabolism phenotyping

The pesticide-degrading isolates IIL-Cl25, IIL-Lc-16, IIL-Luf18, and IIL-Sp24 differed in their susceptibility to antibiotics. While susceptibility of IIL-Lc16 and IIL-Sp24 to the tested antibiotics was identical, IIL-Cl25 and IIL-Luf18 had clear different responses. IIL-Cl25 was the only one susceptible to oxacillin (1 µg), while IIL-Luf18 was the only susceptible to cefepime (30 µg). IIL-Cl25 and IIL-Luf18 were twice more susceptible to clindamycin (2 µg) than IIL-Lc16 and IIL-Sp24 (Fig. 1B).

A contrasting response among isolates was also observed in relation to their capacity to metabolize carbohydrates. IIL-Cl25 and IIL-Lc16 were able to metabolize 28 out of the 49 carbohydrates tested, with 25 of them in common. IIL-Cl25 metabolized D-sorbitol, D-raffinose, and D-turanose, while IIL-Lc16 D-arabinose, L-rhamnose, and potassium gluconate. IIL-Sp24 metabolized 22 of the tested carbohydrates and was the only one to metabolize methyl-α-D-mannopyranoside and glycogen as carbon sources. IIL-Luf18 metabolized only 14 carbohydrates, but it was the only isolate to use potassium 5-ketogluconate as a carbon source (Fig. 1B, Supplementary Figure S3).

#### Genomes assembly and annotation

The whole genome sequencing of isolates IIL-Cl25, IIL-Luf18, IIL-Sp24, and IIL-SusEm yielded from 826 to 1614 Mb assembled nucleotides/genome, with draft genomes of nearly 3 Mb (38% GC) in size, and coverage ranging from 277x to 541x. Quality read filtering recovered 98-99% of the total reads, of which 67-95% were paired reads. The isolate IIL-Lc16 was removed from our analysis due to inconsistencies in the obtained assembly. The number of contigs per draft genome ranged from 129 to 230 contigs (Supplementary Table S9). Among all detected CDS, 66% were functionally annotated and 33-35% were further assigned to EC numbers, 27-28% to GO terms, and 23-24% to KEGG pathways (Fig. 7). Hydrolases (139) and transferases (223) were the major groups of functionally annotated enzymes followed by oxidoreductases (77), ligases (62), lyases (46), and isomerases (44) (Supplementary Table S10).

**Fig. 7.**
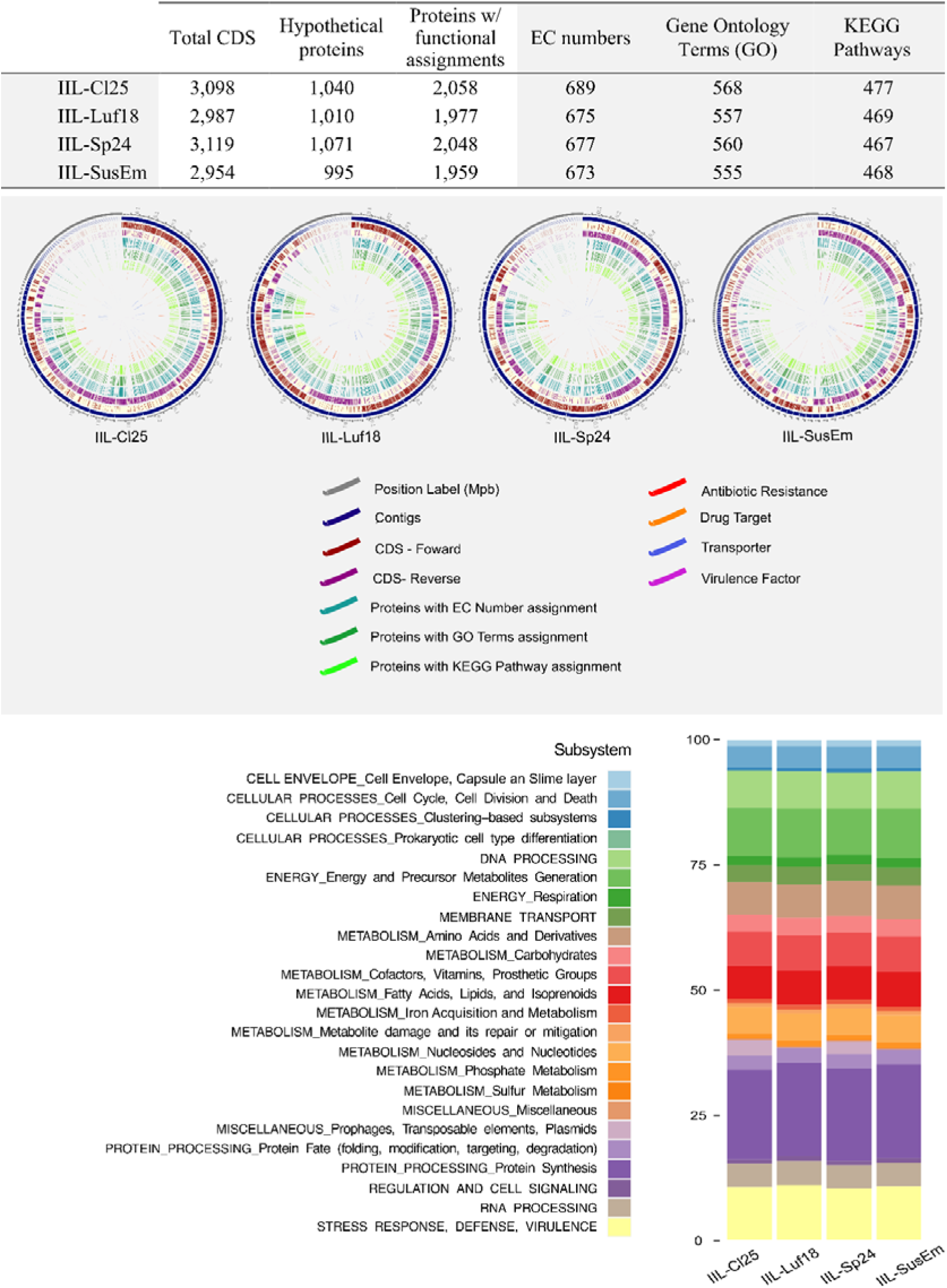
Genome annotation of IIL-Cl25, IIL-Sp24, IIL-Luf18 and IIL-SusEm by Rapid Annotation Subsystem Technology toolkit at BV-BRC website.

Functionally annotated proteins were mapped to KEGG pathways mostly related with the *metabolism of carbohydrates* (23%) (Fig. 8). The proteins mapped to *xenobiotics biodegradation* and *metabolism* pathways represented a diverse range of enzymes, mainly represented by dehalogenases, dehydrogenases, transferases, phosphohydrolases, esterases, deaminases, monooxygenases, and decarboxylases, among others (Fig. 8).

**Fig. 8.**
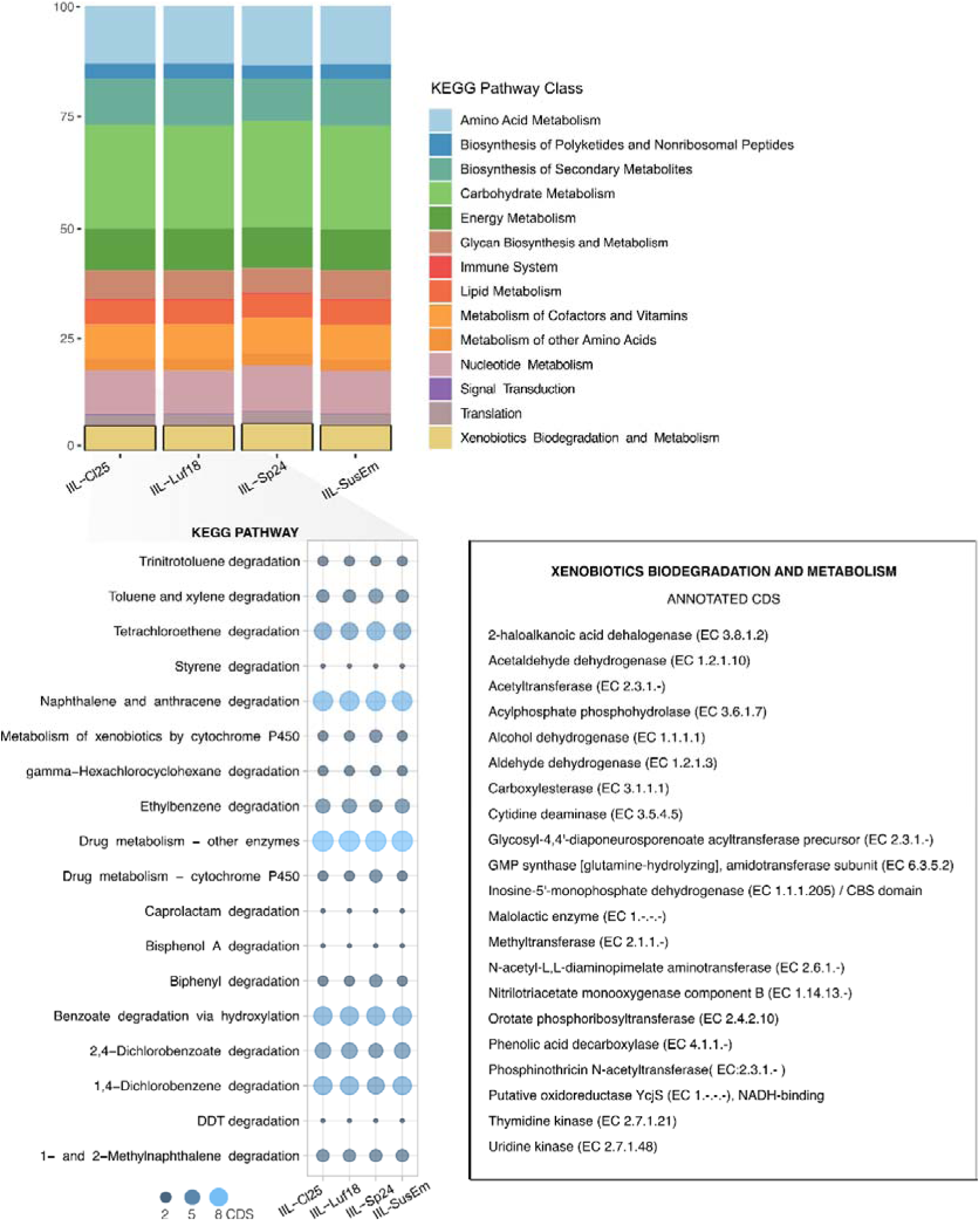
KEGG pathways annotated in the genomes of IIL-Cl25, IIL-Sp24, IIL-Luf18 and IIL-SusEm. Highlighted, genomic sequences mapped with pathways related to xenobiotics biodegradation and metabolism.

A total of 1,605 common coding sequence regions were found in the sequenced isolates (Fig. 9), and 93% of them were functionally annotated. IIL-Sp24 was the most distinctive isolate, with 81 unique proteins, followed by IIL-Cl25 with 25 unique proteins. IIL-Cl25 and IIL-Sp24 shared 37 proteins only between them. IIL-Cl25 still shared other 33 proteins only with IIL-Luf18 and IIL-SusEm. All isolates obtained from insecticide-resistant *S. frugiperda* lines (IIL-Cl25, IIL-Sp24 and IIL-Luf18) shared 18 CDS that were not represented in the genome of the isolate obtained from the insecticide-susceptible reference line (IIL-SusEm). These CDS code the following proteins: *helix*-*turn-helix transcriptional regulator*, *type II toxin-antitoxin system RelB/DinJ family antitoxin*, *bacteriocin immunity pro*tein, *tetracycline resistance gene (TetS)* and transporters from TCDB families *type iv secretory pathway (ivsp), mannose-fructose-sorbose (man),* and *ompa-ompf porin (oop)*.

**Fig. 9.**
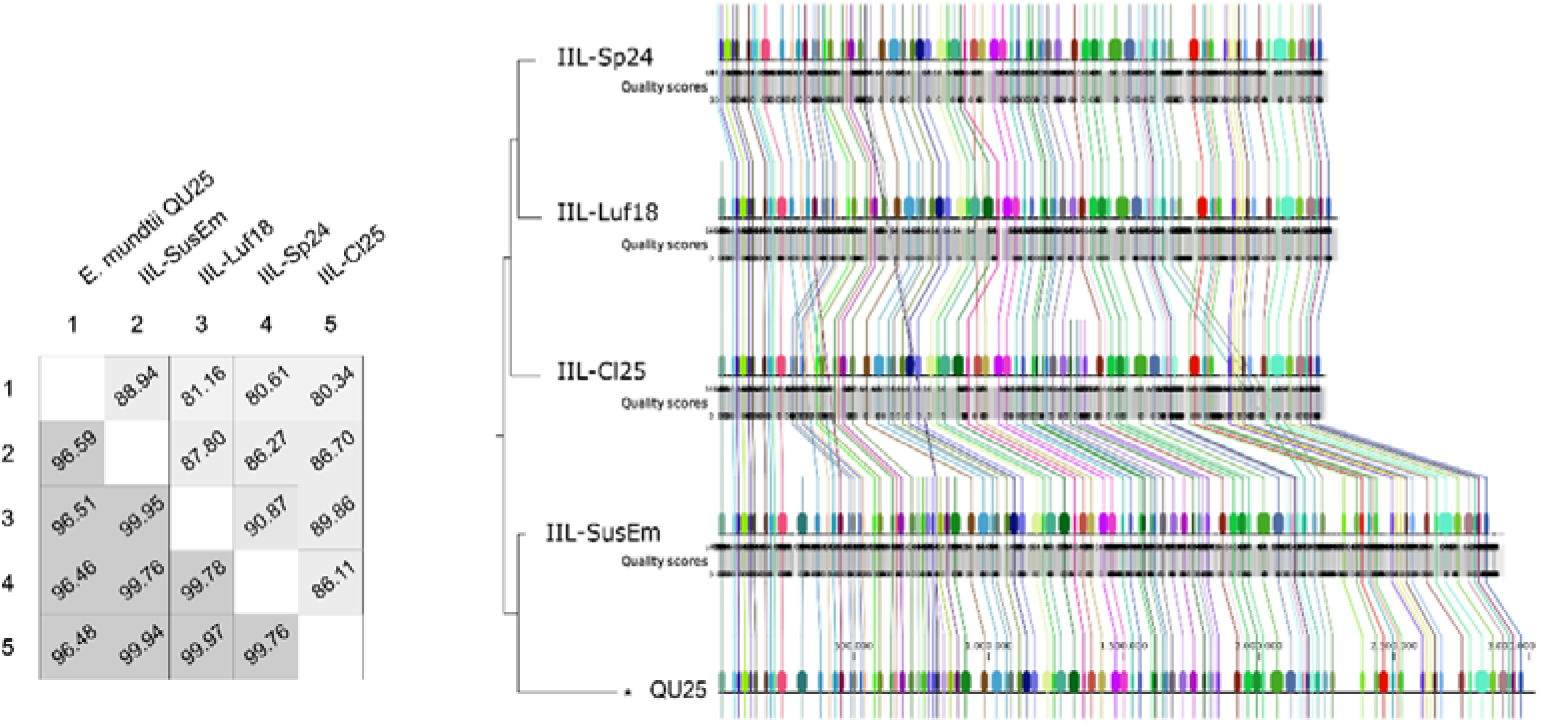
Shared coding regions sequences (CDS) between the isolates IIL-Cl25, IIL-Sp24, IIL-Luf18 and IIL-SusEm. Classification and average amino acid string length of each annotated CDS.

Antibiotic resistant genes common to all isolates were predicted as antibiotic target in susceptible species, antibiotic target protection proteins, antibiotic target modifying enzymes, antibiotic inactivation enzymes, among others (Supplementary Table S11). For CDS predicted as drug targets, stands out the phage-related proteins NinB and lysozyme R (EC 3.2.1.17), annotated only for IIL-Sp24 and IIL-Cl25. All the others predicted CDS were shared among the isolates (Supplementary Table S11).

Coding regions sequences for the TCDB families H (+)- or Na (+)-translocating f-type, v-type and a-type atpase (f-atpase), K (+) transporter (trk) and monovalent cation:proton antiporter-2 (cpa2) were predicted for all sequenced isolates (Supplementary Table S11). Most transporters from type iv secretory pathway (ivsp) and mannose-fructose-sorbose (man) were also shared. Nickase (3.A.7.14.1) from ivsp family and the ABC transporter 3.A.1.5.11, however, were annotated only for IIL-Sp24, while Paraquat-inducible protein B (9.A.69.1.1) was detected only in IIL-Cl25. The phage-related transporters endopeptidase Rz (1.M.1.1.1) from Rz (1) family and the holin/antiholin component S (1.E.2.1.1) were detected only in IIL-Cl25 and IIL-Sp24. The multidrug resistance efflux pump PmrA, as well as the multidrug-efflux transporter transcription regulator BltR were annotated in the genome of all sequenced isolates.

No unique virulence factors were predicted for the isolates analyzed. They all shared the predicted virulence factors (enzymes) *maltose operon transcriptional repressor MalR* - related to biofilm production, *thymidylate synthase* (EC 2.1.1.45), *methionine aminopeptidase* (EC 3.4.11.18), *adenylosuccinate lyase* (EC 4.3.2.2), *ATP-dependent Clp protease* (EC 3.4.21.92), and the *peroxide stress regulator PerR*. Plasmid’s annotation identified mainly hypothetical proteins (Supplementary Table S12), with *tetracycline resistance gene (TetS)* being predicted for all of them.

BGCs searches in AntiSMASH for IIL-SusEm, IIL-Luf18, IIL-Sp24, and IIL-Cl25 genomes detected 2 clusters related to the biosynthesis of secondary metabolites (Supplementary Figure S4). Terpene BGC was detected in all isolates, sharing 36 (IIL-Cl25 and IIL-Luf18) or 46% (IIL-Sp24 and IIL-SusEm) of similarity with the carotenoid BGC from BGC0000647. The second cluster was represented by T3PKS BGC and it shared 23% similarity with the viguiepinol from BGC0000286. This cluster was also detected in all isolates from insecticide-resistant larvae.

#### Whole genome alignment, ANI and dDDH analyses

Genome pairwise alignments of isolates IIL-Cl25, IIL-Luf18, IIL-Sp24, and IIL-Sus-Em ranged from 86 to 90%, with ANI values higher than 99.7%. Lower genome alignments were obtained in pairwise alignments against the reference genome (∼80%), except for IIL-Sus-Em (88.9%) (Fig. 10). Pairwise ANI values of isolates IIL-Cl25, IIL-Luf18, IIL-Sp24, and IIL-Sus-Em against the *E. mundtii* Qu25 reference genome was close to 96.5%. Genome clustering grouped isolates obtained from insecticide-resistant lines of *S. frugiperda* in a cluster separated from that obtained from susceptible larvae, which resolved in a clade containing the reference genome (Fig. 10). Pairwise dDDH analysis agreed with the ANI values obtained for pairwise comparisons between the isolates sequenced and the reference *E. mundtii* strain (dDDH value < 70%), indicating that none of the investigated isolates belong to known species (Fig. 6 and Supplementary Table S8). Therefore, the fragment of the 16S-rDNA previously used for the putative identification of the isolates obtained from *S. frugiperda* [28] did not provide enough information for correct species identification and phylogenetic placement of the studied isolates. We thus propose the isolates obtained from *S. frugiperda* that shared 16S-rDNA sequences identical to those of *E. mundtii* represent a new species based on the ANI and dDDH values obtained, and we propose to name it *Enterococcus spodopteracolus* n. sp., electing the isolate IIL-Cl25 as the genome type species.

**Fig. 10.**
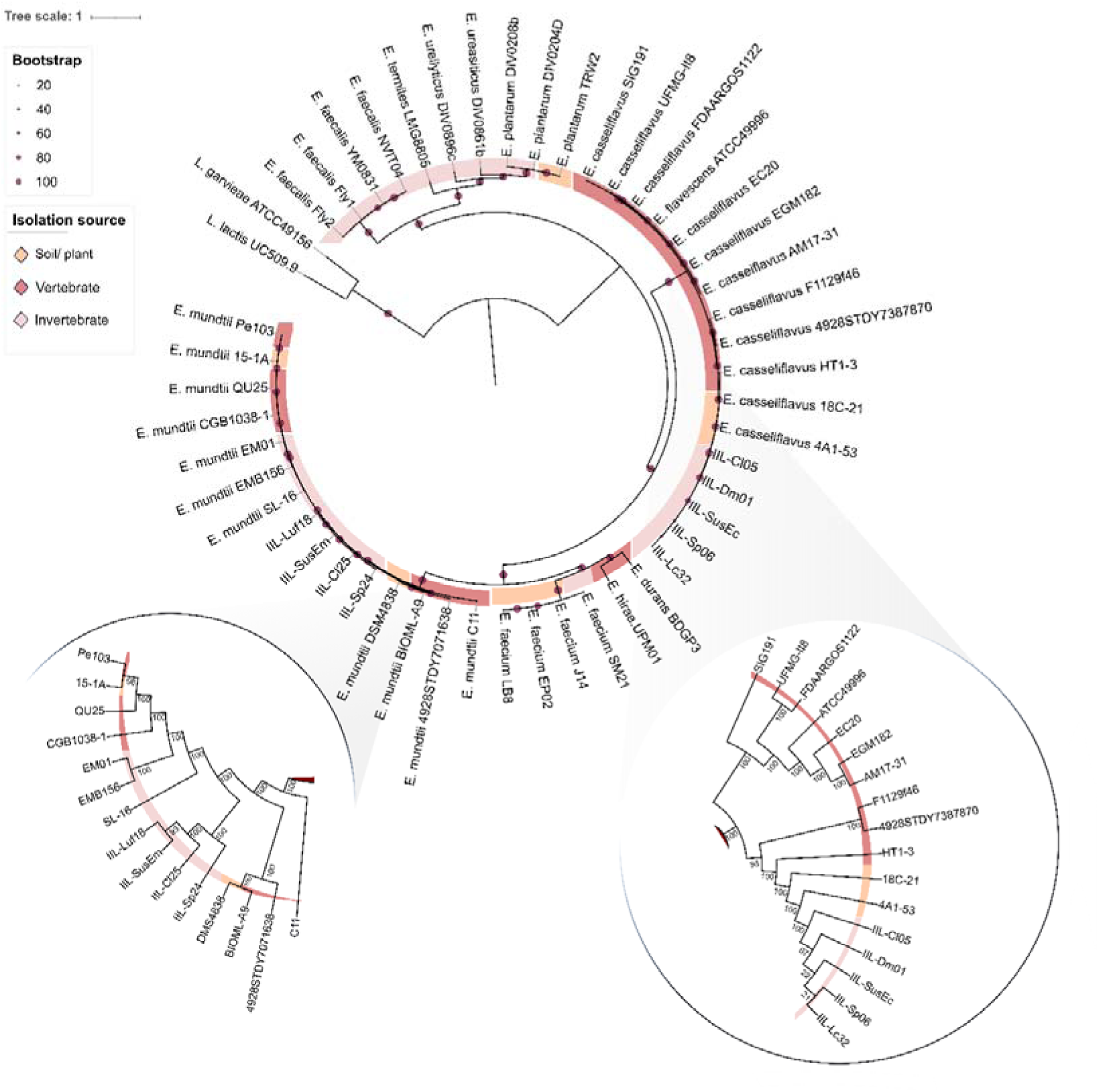
Whole genomes alignment performed at CLC Genomics Workbench (QIAGEN) platform. Percentage of similarity between the isolates IIL-Cl25, IIL-Sp24, IIL-Luf18 and IIL-SusEm, and the reference Qu25 (GenBank: AP013036.1). In the figure, percentage of alignment (AP) (upper) and the average nucleotide identity (ANI) (down).

#### Molecular markers for species differentiation

All six genes that are commonly used as markers for the multilocus sequencing typing of several *Enterococcus* species were highly informative for species differentiation of *E. entomosocium* n. sp. from *E. casseliflavus* and other *Enterococcus* species (Supplementary Figure S5). In the case of *E. spodopteracolus* n. sp*, Gdh* and *PurK* did not provide enough information for species differentiation from *E. mundtii* (Supplementary Figure S5). No differences were observed among the *Enterococcus entomosocium* n. sp. (IIL-CL05, IIL-Dm01, IIL-Lc32, IIL-Sp06, IIL-SusEc) isolates regarding the representative regions of *atpA* (556pb), *ddl* (465bp), *gdh* (530bp), *pstS* (584bp), *purK* (489bp), and *xpt* (456 bp). IIL-SP24 differed from the other *E. spodopteracolus* n. sp. isolates only by a single nucleotide polymorphism (C94A) that results in a non-synonymous mutation (P32T) in the *xpt* gene. The remaining regions analyzed for this species turned out to be identical.

### Phylogenetic analysis

The evolutionary history and relationship of the bacterial isolates from the larval gut of *S. frugiperda* and 28 *Enterococcus* genomes resolved the isolates from *S. frugiperda* in single clades (Fig. 6). The isolates IIL-Cl05, IIL-Sp06, IIL-Lc32 and IIL-SusEc of *Enterococcus entomosocium* n. sp. resolved in a highly supported internal clade, separated from that containing *E. casseliflavus* EC20 and two other *E. casseliflavus* genomes with 100% bootstrap support value (Fig. 6). The highly supported subclade containing the *Enterococcus spodopteracolus* n. sp. IIL-CL25, IIL-Luf18, IIL-Sp24, and IIL-Sus-Em isolates shared a clade with another subclade formed by *E. mundtii* isolates. These subclades are well-supported and are well-separated from other *Enterococcus* species by 100% bootstrap value. Yet, *E. spodopteracolus* n. sp. isolates branched out forming three internal clades with very high bootstrap support values. The first clade to branch out was formed by isolate IIL-Sp24 (100% bootstrap value), the second by IIL-SusEm (100% bootstrap value), and the third by isolates IIL-Luf18 and IIL-Cl25 (86% bootstrap value), all of them with very short branches (Fig. 6).

Our phylogenetic analysis for the determination of habitat specific associations using several species of *Enterococcus* isolated from vertebrates, invertebrates, and soil/plants clustered the gut isolates of *S. frugiperda* in the clades with their closely related species, *E. casseliflavus* and *E. mundtii* (Fig. 11). The gut isolates of *E. entomosocium* n. sp. resolved in the well-separated, most internal clade represented by samples obtained from invertebrates, followed by an intermediary clade with *E. casseliflavus* from soil/plant samples, and the most external clade containing *E. casseliflavus* obtained from vertebrates (Fig. 11).

**Fig. 11.**
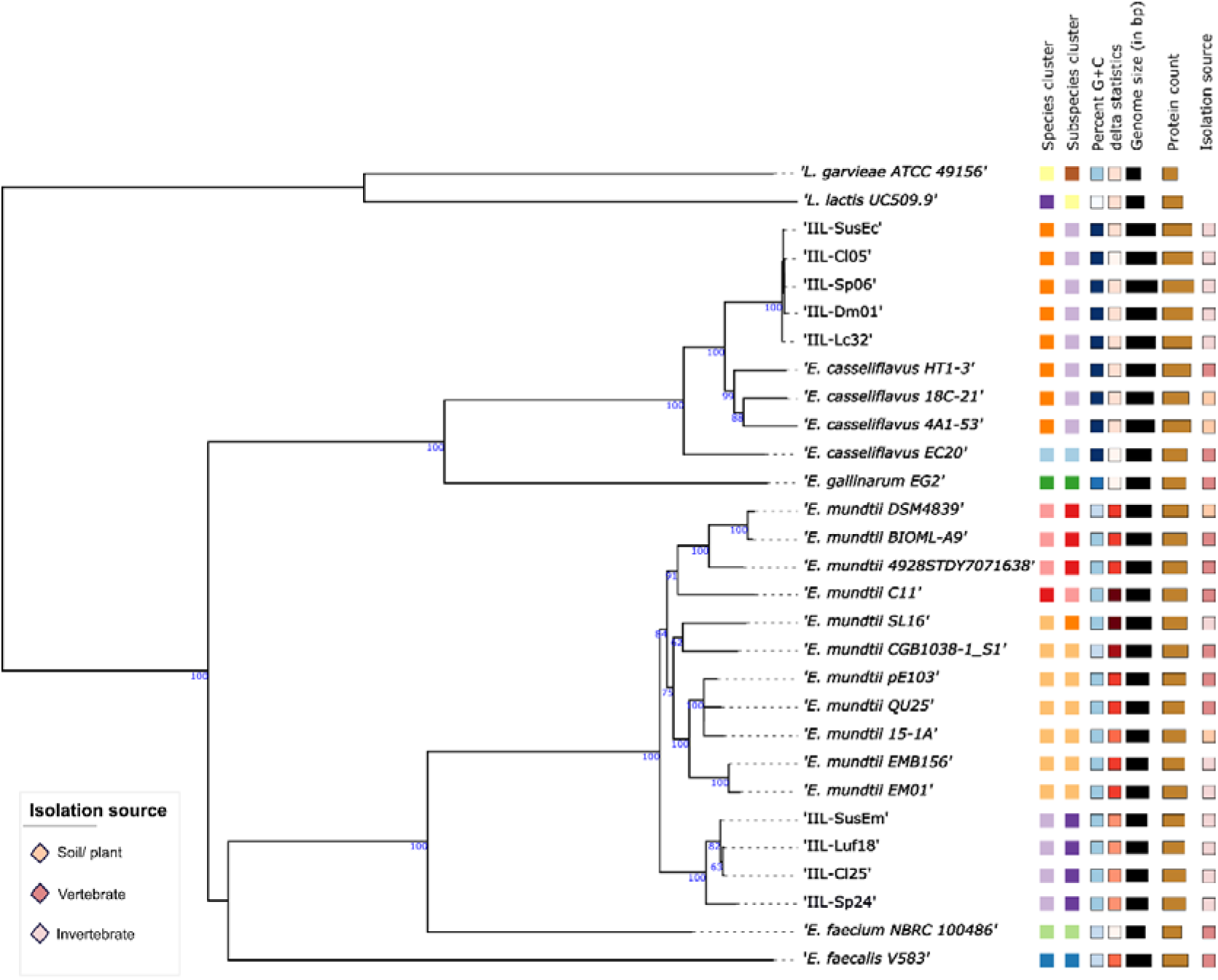
Phylogenetic analysis of *Enterococcus* species collected from different environmental sources, including *Enterococcus entomosocium* n. sp. (IIL-Cl05, IIL-Sp06, IIL-Lc32, and IIL-SusEc) and *Enterococcus spodopteracolus* n. sp. (IIL-Cl25, IIL-Sp24, IIL-Luf18, and IIL-SusEm) from resistant and susceptible strains of *Spodoptera frugiperda.* The phylogenetic tree arrangements with the closest species of *Enterococcus entomosocium* n. sp. (IIL-Cl05, IIL-Sp06, IIL-Lc32, and IIL-SusEc) and *Enterococcus spodopteracolus* n. sp. (IIL-Cl25, IIL-Sp24, IIL-Luf18, and IIL-SusEm) are highlighted. The values in the different branches correspond to bootstrap values. The genomes of *Lactobacillus garvieae* ATCC49156 (GenBank AP009332.1) and *Lactobacillus lactis* UC509.9 (GenBank CP003157.1) were used as external groups in this analysis.

A similar trend was observed for the gut isolates of *E. spodopteracolus* n. sp. A much higher variation was observed for *E. mundtii*, the closely related species of *E. spodopteracolus* n. sp. The isolates of *E. spodopteracolus* n. sp. clustered altogether in a clade close to three isolates of *E. mundtii* (SL-16, EMB156, and EM01) obtained from invertebrates. *Enterococcus spodopteracolus* n. sp. and *E. mundtii* SL-16, EMB156, and EM01 are all associated with invertebrates, and all resolved in between external clusters with *E. mundtii* isolates associated with vertebrates and soil/plant samples (Fig. 11).

The clustering obtained prompted us to investigate the genomic similarities among the several isolates of *E. entomosocium* n. sp. and *E. spodopteracolus* n. sp. with isolates of *E. casseliflavus* and *E. mundtii* from different environmental sources. The whole-proteome-based phylogeny corroborated the position of the *Enterococcus* symbionts of *S. frugiperda* as two new species (Fig. 12 and Supplementary Table S13), but also provided additional information for the correct systematic positioning of isolates currently reported as *E. mundtii* and *E. casseliflavus*. This analysis separated all representatives of *E. mundtii* in three different species cluster according to dDDH value (Fig. 12 and Supplementary Table S13), all these clusters are distinct from the species cluster formed with the several isolates of *E. spodopteracolus* n. sp. The species cluster contained the isolates *E. mundtii* DSM4839, *E. mundtii* BIOML-A9, and *E. mundtii* 4928SDTY7071638 belong to the *E. mundtii* type species group. The isolate *E. mundtii* C11 formed represents alone a species cluster. The third species cluster contains seven other isolates, with *E. mundtii* SL16 belonging to a different subspecies cluster. All *E. entomosocium* n. sp. isolates joined in a single species cluster with two isolates from soil/plants (18C-21 and 4A1-53) and one isolate from vertebrates (HT1-3). This species cluster differed from the species cluster in which the type *E. casseliflavus* EC20 resolved. Comparative genomic analysis of isolates HT1-3 (Genbank VODL00000000), 18C-21 (Genbank JAAOLS000000000) and 4A1-53 (Genbank JAAMSH000000000.1) previously reported as *E. casseliflavus* demonstrated that these isolates in fact belong to the same species cluster of *E. entomosocium* n. sp. (dDDH>70%) (Fig. 12 and Supplementary Table S13).

**Fig. 12.** Phylogenetic analysis between isolates of *Enterococcus entomosocium* n. sp. (IIL-Cl05, IIL-Sp06, IIL-Lc32, and IIL-SusEc) and *Enterococcus spodopteracolus* n. sp. (IIL-Cl25, IIL-Sp24, IIL-Luf18, and IIL-SusEm) from resistant and susceptible strains of *Spodoptera frugiperda* and the closest species, *Enterococcus mundtii* and *Enterococcus casseliflavus*. Whole-proteome-based phylogeny performed at the Type (Strain) Genome Server (TYGS). The cut-offs of 70% and 79% for digital DNA: DNA hybridization (dDDH) value were adopted for, respectively, species and subspecies delineation. The values in the different branches correspond to bootstrap values.

## Discussion

### Genotypic and phenotypic diversity

Isolates of *Enterococcus entomosocium* n. sp. and *Enterococcus spodopteracolus* n. sp. carried high genotypic similarities within each species (Fig. 5 and 10, Supplementary Table S8). The phenotypic plasticity observed among *E. spodopteracolus* isolates (Fig. 1b) cannot be explained by variations in their genomes, suggesting that their plasticity in degrading different classes of pesticides [28], responding to antibiotics and metabolizing different sources of sugars is due to epigenetic mechanisms [61, 62]. In a range of bacterial species, such as *Escherichia coli* [63–65], *Streptococcus pneumoniae* [66–68] and *Helicobacter pylori* [69, 70], DNA methylation control phase variation of gene expression, which may generate phenotypic cell variants without genetic mutation. Further studies, however, are necessary to support this hypothesis.

### Potential on insecticides metabolization

The diverse enzymatic repertoire available in microorganisms and their remarkable catabolic potential make them key players in detoxification processes [71, 72]. The outstanding variability of enzymes acting on natural and synthetic chemistries in bacteria allows their exploitation for bioremediation of contaminated areas [71, 73–75], and their selection by host insects that are benefitted by the bacterial metabolization of xenobiotics [28, 76, 77]. Hydrolases of insect-associated bacteria were already demonstrated to be involved in the reduction of organophosphate toxicity in the stink bug *Riptortus pedestris* (Hemiptera: Alydidae) [78] and in the fruit fly *Bactrocera dorsalis* (Diptera: Tephritidae) [79].

Our genomic analyses of the isolates of *Enterococcus* obtained from insecticide-resistant larvae of *S. frugiperda* [28] demonstrated they are well served of an enzymatic machinery for xenobiotic degradation (Fig. 3 and 8, Supplementary Tables S5 and S10), which is represented by several carboxylesterases, dehalogenases and dehydrogenases that participate in the transformation and/or breakdown of insecticides [80–82]. Hydrolases acting on halide, acid anhydrides and carbon-nitrogen bonds, and transferases, oxidoreductases, and carboxylases are also part of the enzymatic repertoire that act on xenobiotic metabolization, and they were detected in the *Enterococcus* isolates studied (Supplementary Tables S5 and S10). These enzymes are associated with drug metabolism in a range of bacterial genera such as *Pseudomonas, Bacillus, Microbacterium*, and *Arthrobacter* [80, 83, 84].

The enzymatic machinery identified in all *S. frugiperda*-associated *Enterococcus* isolates we investigated is equipped with bacterial enzymes that have been shown to degrade the organic chemistries they were selected upon, such as pyrethroids (lambda-cyhalothrin and deltamethrin) [85], benzoyphenylureas (lufenuron) [86], and spinosyns (spinosad) [87]. Despite the sequenced isolates of *Enterococcus* associated with susceptible and insecticide-resistant larvae of *S. frugiperda* carry several enzymes involved in xenobiotic degradation, the efficiency of microbial detoxification of several pesticides, such as teflubenzuron and chlorpyrifos has been much higher when exposed to microbial consortia [88, 89]. The investigated isolates of *Enterococcus* certainly played a role in the detoxification of pesticides in the gut of the *S. frugiperda* strain they were originally selected from [28], but they are not the only ones capable to degrade insecticides that were selected from the gut of insecticide resistant lines of *S. frugiperdae* [28]. Thus, the evaluation of the contribution of gut bacteria in insecticide degradation in host insects, such as *S. frugiperda* should consider the potential contribution of the bacteria consortium for a reliable evaluation of the full potential for detoxification of the gut microbiota.

Our analysis also supports the original hypothesis of [28] that gut bacteria are also exposed to the selection pressure by insecticides as their host insects are, as we detected differences in the gene content among the *Enterococcus* isolates sequenced, with clear phenotypic differences in the catabolism of carbohydrates by *E*. *spodopteracolus* n. sp. Changes in the genome of microorganisms for adaptation to stress conditions can occur through single carbon catabolite repression mutations, as demonstrated for the mutualistic association of *Escherichia coli* to the stinkbug *Plautia stali* [90], or through genome streamlining as nicely demonstrated for the plant mutualist strains of *Bradyrhizobium diazoefficiens* adapted to different abiotic stressors [91]. But since the enzymatic repertoire involved in *xenobiotic degradation* in isolates capable of degrading insecticides was also observed in the isolates that were unable to grow on insecticide-selective media (isolated from a susceptible *S. frugiperda* line), we could not discard epigenetic mechanisms as a source of selection in insecticide-degrading isolates.

Besides the production of enzymes acting on the modification or degradation of pesticides, the gut microbiota can mediate xenobiotic detoxification through biofilm protection [92]. Biofilm protection by *E. spodopteracolus* n. sp. inhabiting the gut of *S. frugiperda* is indicated by the recognition of the *maltose operon transcriptional repressor MalR,* a *bopD*-encoded sugar-binding transcriptional regulator that participates in biofilm production in *Enterococcus faecalis* [93, 94], and has been annotated in all isolates of *E. spodopteracolus* n. sp. (Supplementary Table S11). Bacterial bioaccumulation has also been demonstrated as a mechanism that can interfere with the interactions of therapeutic drugs with humans [95], an ability earlier shown for *Escherichia coli* to bioaccumulate the pesticide fipronil [73]. Several influx transporters reported in *E. coli* (*atpA*, *atpB, atpD, ssuC*, *ycdG*, and others) [96] have also been annotated in the genome of *E. entomosocium* n. sp. and *E. spodopteracolus* n. sp. Efflux systems are also direct mechanisms involved in xenobiotic detoxification by gut bacteria [92]. Genome analysis of *E. entomosocium* n. sp. and *E. spodopteracolus* n. sp. allowed the identification of several efflux pumps that are known to actively pump several xenobiotics out of bacterial cells, such as the *multidrug resistance efflux pump PmrA* and the *multidrug-efflux transporter transcription regulator BltR* identified in *Streptomyces pneumoniae* and *Bacillus subtilis*, respectively [97–99]. However, we believe efflux pumps play a small role, if any, in xenobiotic detoxification by both enterococci associated with *S. frugiperda,* as these bacteria are mainly free-living inhabitants of the gut lumen.

Bacteria can also contribute to host xenobiotic degradation indirectly by regulating the host defense mechanisms following host exposure to stressors [92], resulting in an alteration of the host immune system [26, 100, 101]. Annotated virulence factors (Supplementary Tables S6 and S11) and phage-related proteins suggest the potential contribution of the *Enterococcus* species studied to host xenobiotic detoxification. Further analysis regarding gene expression would be necessary to confirm the direct and indirect contributions of *E. entomosocium* n. sp. and *E. spodopteracolus* n. sp. to the metabolism of xenobiotics and detoxification in *S. frugiperda* larvae.

### Host interaction - Virulence, infectivity, and functional contribution

The host-microbiota interaction and the ways by which they relate to each other are critical points for the characterization and better understanding of symbiotic relationships [102–104]. Pathogenicity traits related to infection processes were detected in all *E. entomosocium* n. sp. and *E. spodopteracolus* n. sp. genomes, including virulence factors (Supplementary Tables S6 and S11) and enzymes associated with host infection and colonization (Supplementary Tables S5 and S10). The *peroxide stress regulator PerR* (FUR family) and the *ATP-dependent Clp protease proteolytic subunit ClpP* (EC 3.4.21.92) of *E. entomosocium* n. sp. and *E. spodopteracolus* n. sp. were shown to influence the oxidative-stress response and virulence of *E. faecalis* and *Staphylococcus aureus* [105, 106]. The *maltose operon transcriptional repressor* MalR has been linked to bacterial colonization and persistence in the host gut through biofilm formation [93, 94]. Both *ATP-dependent Clp protease proteolytic subunit ClpP* and *maltose operon transcriptional repressor MalR* were also present in the genome of *Enterococcus mundtii* EM01 strain - the pathogen responsible for Flacherie disease in silkworms [107].

The contribution of *Enterococcus* species to the competitive exclusion of pathogenic bacteria in the host gut microbiota by the synthesis and release of bacteriocins has been reported for a range of insect hosts [17, 108]. The bacteriocin mundticin was first identified as a bioactive molecule produced by *Enterococcus mundtii* NFRI 7393 isolated from grass silage [109], and more recently shown to be secreted by *E. mundtii* associated with the larval gut of *Spodoptera littoralis* and proposed to protect the host gut from establishing pathogenic interactions [17]. Mundcitin is also available in the genome of *E. spodopteracolus* n. sp. (Supplementary Table S10), suggesting this bacterium can provide additional defensive mechanisms against pathogen colonization of the gut. As lactic acid bacteria, both *E. entomosocium* n. sp. and *E. spodopteracolus* n. sp. may be able to produce organic acids during growth, reducing the medium pH to facilitate their growth in alkaline environments like the gut of lepidopteran larvae [13, 110].

Insects generally cannot synthetize B vitamins, which are supplied through diet or by microbial symbionts [111]. Enzymes related to pyridoxin (vitamin B6) biosynthesis were annotated in all *E. entomosocium* isolates (Supplementary Table S5), indicating a potential nutritional contribution of *E. entomosocium* n. sp. to the *S. frugiperda* host. Vitamin B contribution is common for symbionts associated with hematophagous insects, as the tsetse’s obligate symbiont *Wigglesworthia glossinidia* [112]. Key genes related with the biosynthesis of the essential amino acid L-tryptophan – *3*ℒ*phosphoshikimate 1*ℒ*carboxyvinyltransferase (aroA), shikimate kinase (aroK), phosphoribosylanthranilate isomerase (TrpF),* and *anthranilate synthase (trpG)* - were detected in *E. entomosocium* n. sp. and *E. spodopteracolus* n. sp. genomes (Supplementary Tables S5 and S10). Genes encoding the alpha and beta subunits of the enzyme *tryptophan synthase* (EC 4.2.1.20) were annotated in all *E. entomosocium* n. sp. isolates (Supplementary Table S5). These genes have been associated with L-tryptophan biosynthesis in *E. casseliflavus* ECB140 [16].

### Systematic positioning symbionts

16S rRNA sequencing has been shown to be a reliable molecular marker for bacterial species identification, particularly for species of clinical importance [113]. But 16S rRNA gene sequence not always lead to correct species identification, even when identical sequences are obtained [114], which prompt the proposition and development of polyphasic taxonomy [115]. Polyphasic taxonomy has been later challenged based on the large number of species to be described and the extensive number of phenotypic assays required in defense of a genome-based taxonomy [116], leading to the development of minimal standards for the use of genomic data in prokaryote taxonomy [117].

Average nucleotide identity (ANI) and digital DNA: DNA hybridization (dDDH) have been described as the most acceptable and used standards for delineating new species through whole genome sequence data [117]. ANI values of 95∼96 and dDDH of 70% were proposed and validated as cut-off points for delimiting new species [53, 54], fully endorsing our propositions of the two new species (Fig. 5 and 10, Supplementary Table 8), *Enterococcus entomosocium* n. sp. and *Enterococcus spodopteracolus* n. sp. as gut symbionts of the lepidopteran pest *Spodoptera frugiperda,* and our call for a reappraisal of the *Enterococcus* species associated with insects.

## Conclusions

Comparative genomics analysis allowed us to better characterize the enterococci symbionts of *S. frugiperda* with the recognition of two new species of enterococci that were named *Enterococcus entomosocium* n. sp. and *Enterococcus spodopteracolus* n. sp. Both species present features on their genomes that indicate their potential contribution to host detoxification, as well as that support the hypothesis that gut bacteria are also exposed to insecticide selection pressure, as their host insects are.

## Author Contributions

Conceptualization: AFG, LGA and FLC; Data curation: AFG and LGA; Formal analysis: AFG, LGA and FLC; Funding acquisition: FLC; Investigation: AFG (comparative genomics) and LGA (phenotyping); Methodology: AFG, LGA and FLC; Project administration: FLC; Supervision: FLC; Writing original draft: AFG and FLC; Writing, review & editing: all authors read and approved the manuscript.

## Data Availability

The datasets generated and/analyzed during this study will be available at the National Center for Biotechnology Information (https://www.ncbi.nlm.nih.gov/) under the bioprojects PRJNA943201(accession numbers SAMN33713714-SAMN33713718) and PRJNA944113 (accession numbers SAMN33735643-SAMN33735646).

---

**Figure S1.**
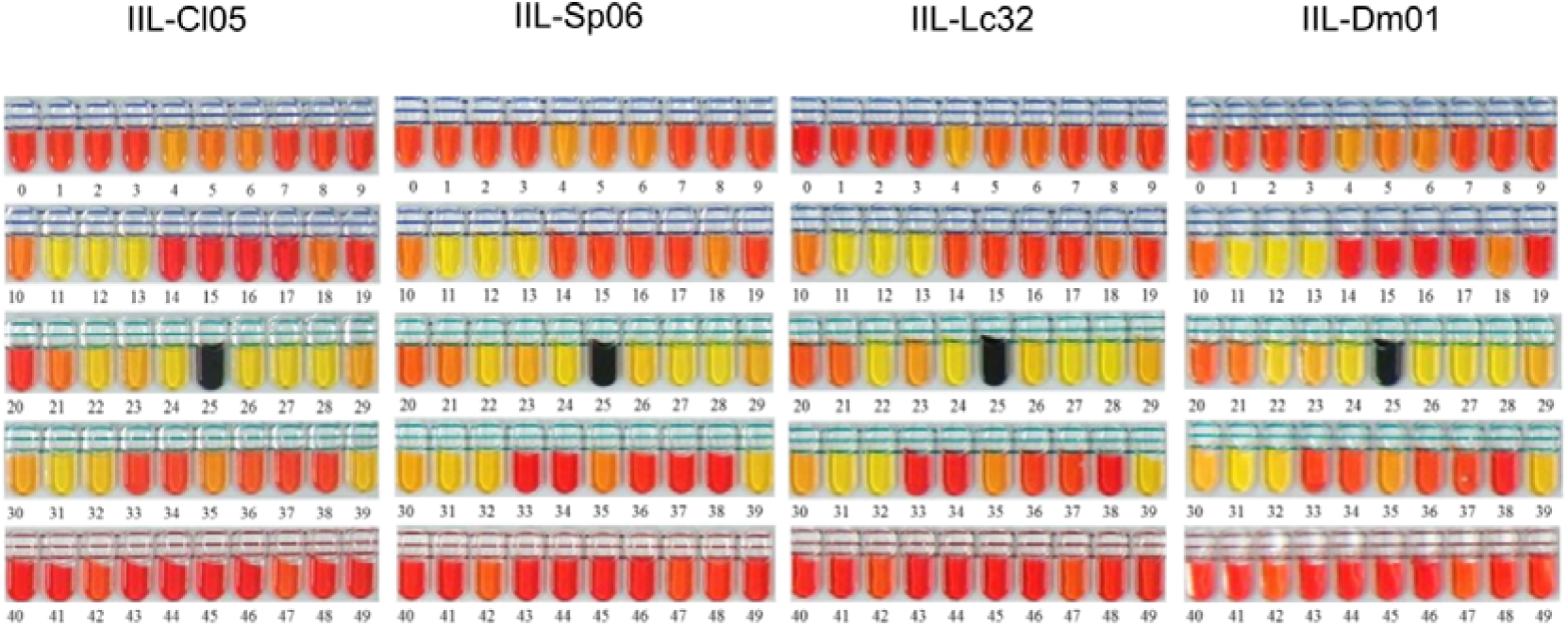
Carbohydrate metabolism of isolates IIL-Cl05, IIL-Sp06, IIL-Lc32 and IIL-Dm01 of *Enterococcus entomosocium* n. sp. API 50 CH galleries (BioMérieux, France) after 48 hours of incubation. Composition of galleries: 0 Control; 1 Glycerol; 2 Erythritol; 3 D-arabinose; 4 L-arabinose; 5 D-ribose; 6 D-xylose; 7 L-xylose; 8 D-adonitol; 9 Methyl-ßd-xylopyranoside; 10 D-galactose; 11 D-glucose; 12 D-fructose; 13 D-mannose; 14 L-sorbose; 15 L-rhamnose; 16 Dulcitol; 17 Inositol; 18 D-mannitol; 19 D-sorbitol; 20 Methyl-αd-mannopyranoside; 21 Methyl-αd-glucopyranoside; 22 N-acetylglucosamine; 23 Amygdalin; 24 Arbutin; 25 Esculin ferric citrate; 26 Salicin; 27 D-cellobiose; 28 D-maltose; 29 D-lactose (bovine origin); 30 D-melibiose; 31 D-saccharose (sucrose); 32 D-trehalose; 33 Inulin; 34 D-melezitose; 35 D-raffinose; 36 Amidon (starch); 37 Glycogen; 38 Xylitol; 39 Gentiobiose; 40 D-turanose; 41 D-lyxose; 42 D-tagatose; 43 D-fucose; 44 L-fucose; 45 D-arabitol; 46 L-arabitol; 47 Potassium gluconate; 48 Potassium 2-ketogluconate; 49 Potassium 5-ketogluconate.

**Figure S2.**
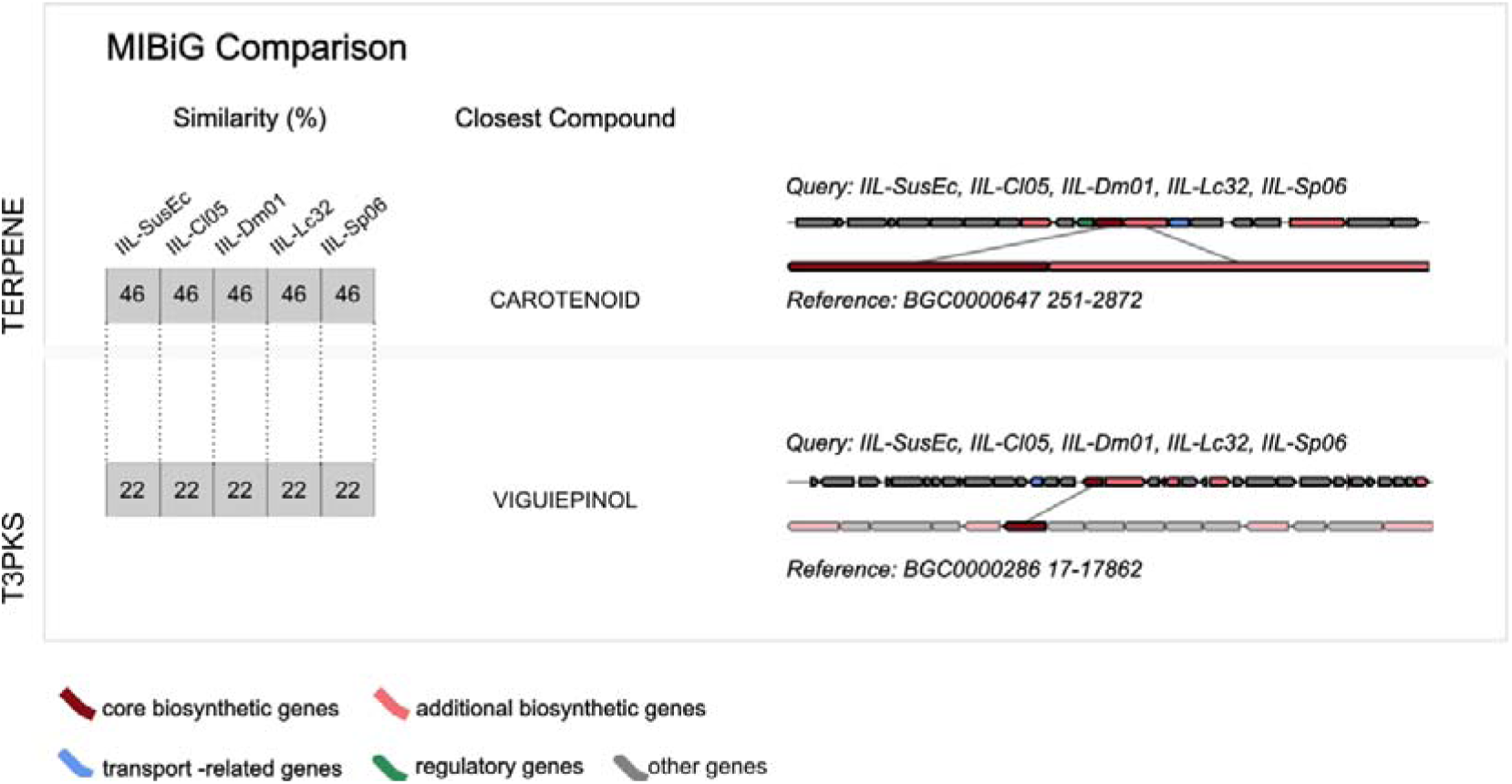
Biosynthetic gene clusters (BGCs) detected by AntiSMASH in the genome of isolates IIL-Sp06, IIL-Dm01, IIL-Cl05, IIL-Lc32 and IIL-SusEc of *Enterococcus entomosocium* n. sp. BGCs type and similarity (percentage of genes) with the closest known compound from MiBIG database.

**Figure S3.**
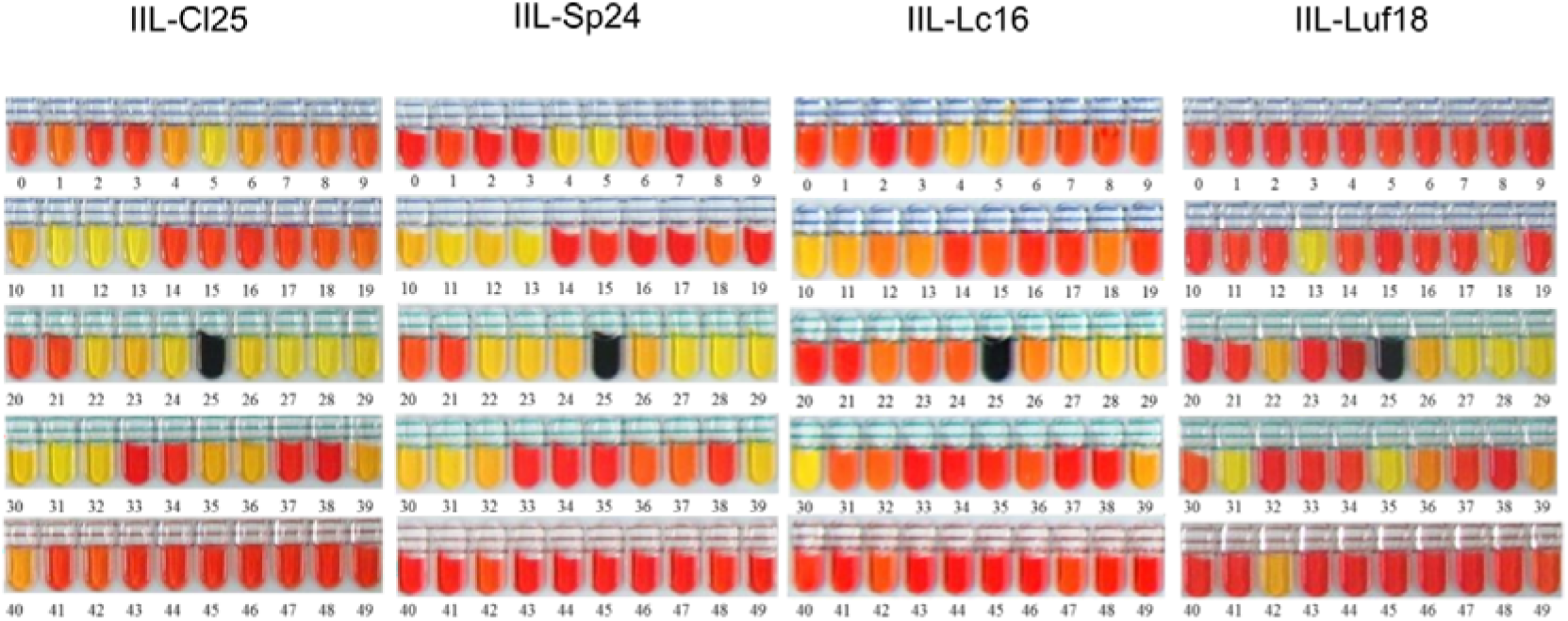
Carbohydrate metabolism of isolates IIL-Cl25, IIL-Sp24, IIL-Lc16 and IIL-Luf18 of of *Enterococcus spodopteracolus* n. sp. API 50 CH galleries (BioMérieux, France) after 48 hours of incubation Composition of galleries: 0 Control; 1 Glycerol; 2 Erythritol; 3 D-arabinose; 4 L-arabinose; 5 D-ribose; 6 D-xylose; 7 L-xylose; 8 D-adonitol; 9 Methyl-ßd-xylopyranoside; 10 D-galactose; 11 D-glucose; 12 D-fructose; 13 D-mannose; 14 L-sorbose; 15 L-rhamnose; 16 Dulcitol; 17 Inositol; 18 D-mannitol; 19 D-sorbitol; 20 Methyl-αd-mannopyranoside; 21 Methyl-αd-glucopyranoside; 22 N-acetylglucosamine; 23 Amygdalin; 24 Arbutin; 25 Esculin ferric citrate; 26 Salicin; 27 D-cellobiose; 28 D-maltose; 29 D-lactose (bovine origin); 30 D-melibiose; 31 D-saccharose (sucrose); 32 D-trehalose; 33 Inulin; 34 D-melezitose; 35 D-raffinose; 36 Amidon (starch); 37 Glycogen; 38 Xylitol; 39 Gentiobiose; 40 D-turanose; 41 D-lyxose; 42 D-tagatose; 43 D-fucose; 44 L-fucose; 45 D-arabitol; 46 L-arabitol; 47 Potassium gluconate; 48 Potassium 2-ketogluconate; 49 Potassium 5-ketogluconate.

**Figure S4.**
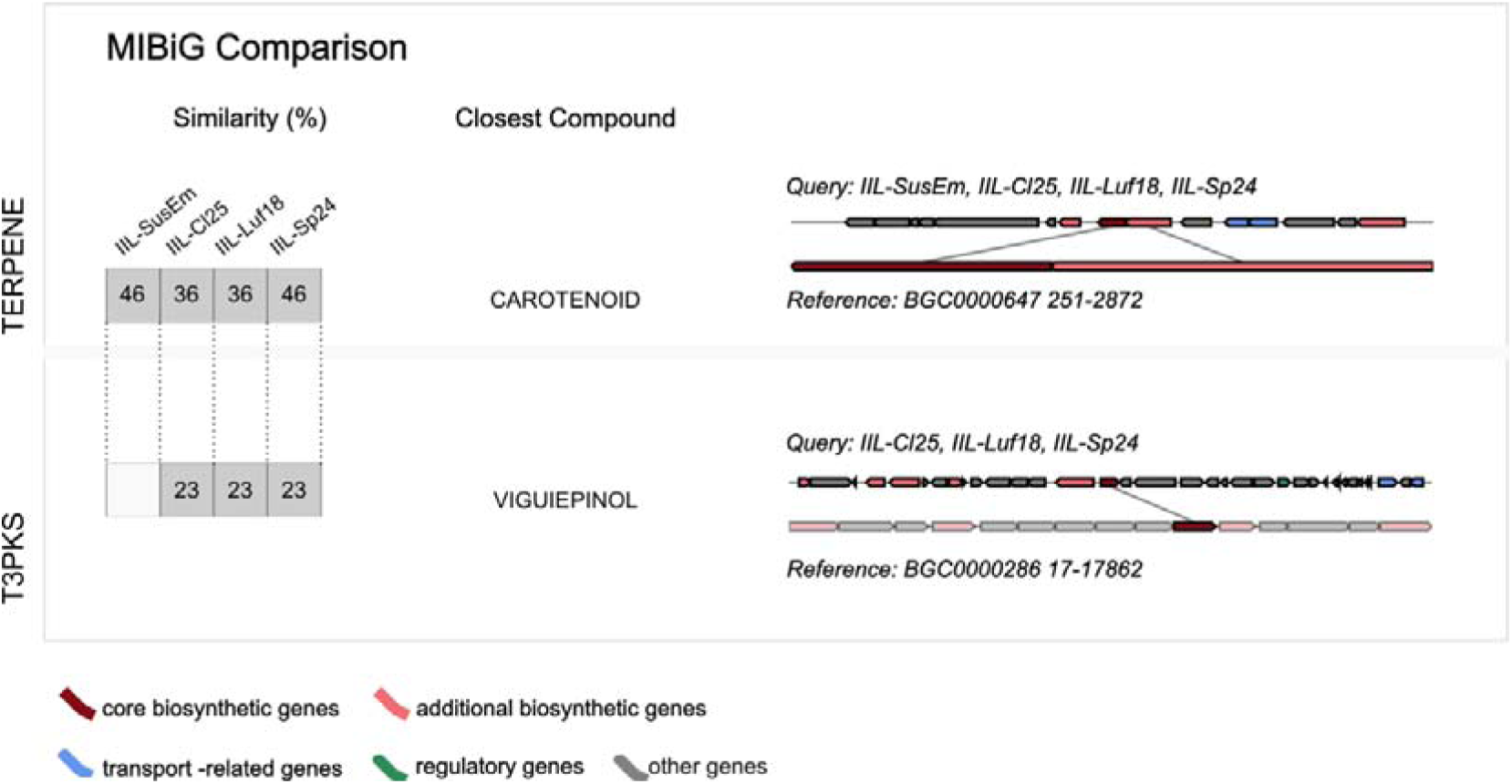
Biosynthetic gene clusters (BGCs) detected by AntiSMASH in the genome of isolates IIL-Sp24, IIL-Cl25, IIL-Luf18 and IIL-SusEm of *Enterococcus spodopteracolus* n. sp. BGCs type and similarity (percentage of genes) with the closest known compound from MiBIG database.

**Figure S5.**
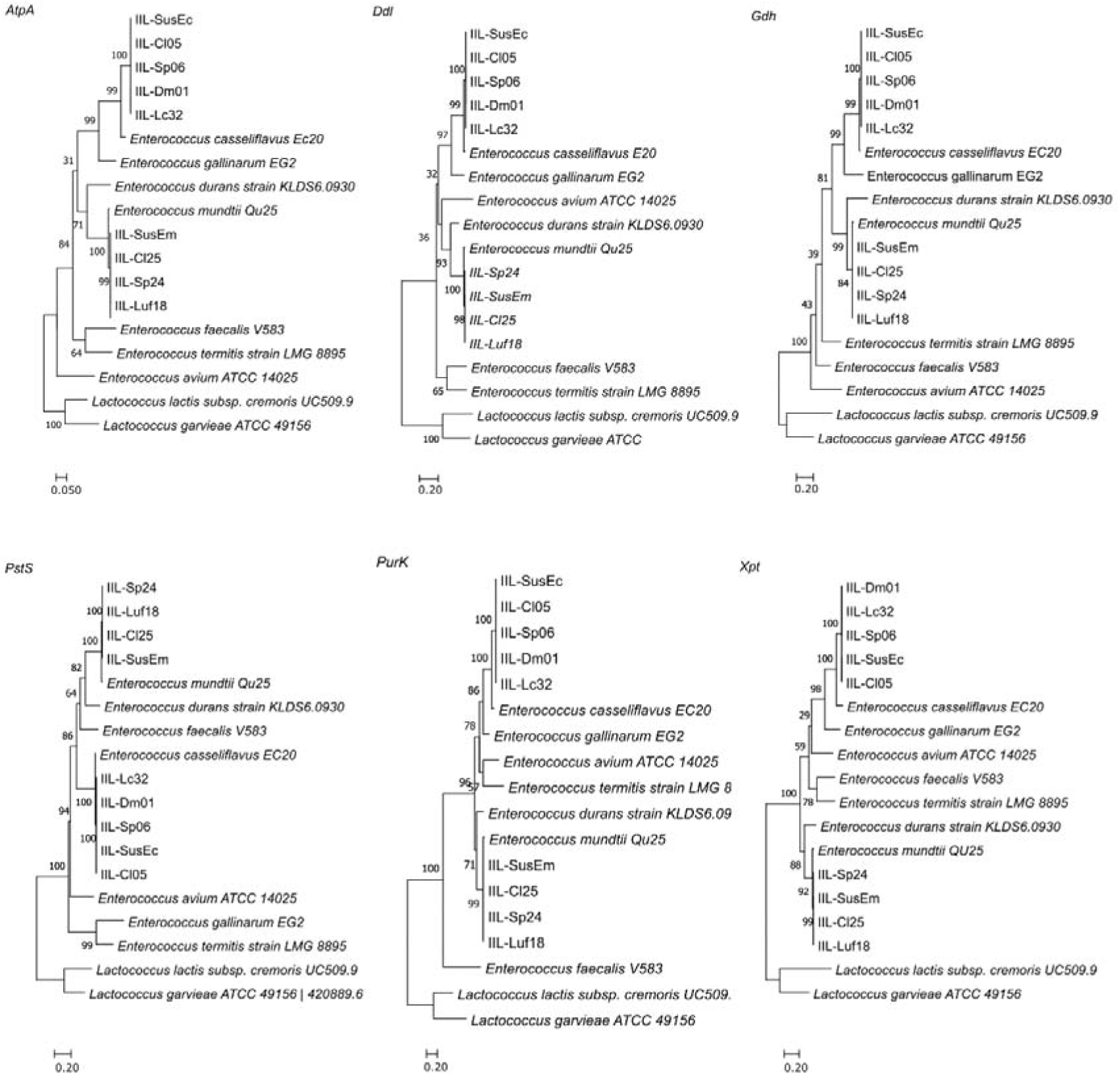
Individual genes phylogenetic analysis showing their potential to discriminate of *Enterococcus entomosocium* n. sp (IIL-Cl05, IIL-Dm01, IIL-Lc32, IIL-SusEc) and, for most of them, to separate of *Enterococcus spodopteracolus* n. sp. (IIL-Cl25, IIL-Luf18, IIL-Sp24, IIL-SusEm) from other *Enterococcus* species. Both *gdh* and *purK* genes are not good candidates for discriminating *E. frugiperdae* from *E. mundtii* Qu25 (84% and 99% bootstrap value, respectively). The values in the different branches correspond to bootstrap values. The strains *Enterococcus avium* ATCC 14025 (GenBank ASWL00000000), *Enterococcus durans* KLDS6.0930 (GenBank CP012384.1), *Enterococcus casseliflavus* EC20 (GenBank CP004856.1), *Enterococcus faecalis* V583 (GenBank AE016830.1), *Enterococcus gallinarum* EG2 (GenBank ACAJ00000000.1), *Enterococcus mundtii* Qu25 (GenBank AP013036.1) and *Enterococcus termites* LMG 8895 (GenBank MIJY00000000.1) were used as reference, and *Lactobacillus garvieae* ATCC49156^T^ (GenBank AP009332.1) and *Lactobacillus lactis* UC509.9^T^ (GenBank CP003157.1) as external groups in this analysis.

## References

1. Icaza-Chávez ME (2013) Gut microbiota in health and disease. Rev Gastroenterol Mex (English Edition) 78:240–248. https://doi.org/10.1016/j.rgmxen.2014.02.009

2. Tacconi E, Palma G, Biase D de, et al (2023) Microbiota effect on trimethylamine N-oxide production: From cancer to fitness—a practical preventing recommendation and therapies. Nutrients 15:563. https://doi.org/10.3390/nu15030563

3. Tsuchida T, Koga R, Horikawa M, et al (2010) Symbiotic bacterium modifies aphid body color. Science 330:1102–1104. https://doi.org/10.1126/science.1195463

4. Schmidt K, Engel P (2021) Mechanisms underlying gut microbiota-host interactions in insects. J Exp Biol 224(Pt 2): jeb207696. https://doi: 10.1242/jeb.207696.

5. Wang G, Berdy BM, Velasquez O, et al (2020) Changes in microbiome confer multigenerational host resistance after sub-toxic pesticide exposure. Cell Host Microbe 27:213–224. https://doi.org/10.1016/j.chom.2020.01.009

6. Zheng H, Perreau J, Powell JE, et al (2019) Division of labor in honey bee gut microbiota for plant polysaccharide digestion. Proc Natl Acad Sci U.S.A 116 (51):25909–25916. https://doi.org/10.1073/pnas.1916224116

7. Engel P, Moran NA (2013) The gut microbiota of insects – diversity in structure and function. FEMS Microbiol Rev 37:699–735. https://doi.org/10.1111/1574-6976.12025

8. Kucuk RA (2020) Gut bacteria in the holometabola: A review of obligate and facultative symbionts. J Pest Sci 20:1–12. https://doi.org/10.1093/jisesa/ieaa084

9. Scoble MJ (1992) The Lepidoptera: Form, function and diversity. OxfordUK: Oxford University Press

10. Wahlberg N, Wheat CW, Peña C (2013) Timing and patterns in the taxonomic diversification of Lepidoptera (butterflies and moths). PLoS One 8:. https://doi.org/10.1371/journal.pone.0080875

11. Holloway JD, Bradley JD, Carter DJ (1987) CIE Guides to insects of importance to man. I. Lepidoptera. CAB International

12. Voirol LRP, Frago E, Kaltenpoth M, et al (2018) Bacterial symbionts in Lepidoptera: Their diversity, transmission, and impact on the host. Front Microbiol 9:556. https://doi.org/10.3389/fmicb.2018.00556

13. Zhang X, Zhang F, Lu X (2022) Diversity and functional roles of the gut microbiota in Lepidopteran insects. Microorganisms 10 (6):1234. https://doi: 10.3390/microorganisms10061234

14. Hammer TJ, Janzen DH, Hallwachs W, et al (2017) Caterpillars lack a resident gut microbiome. Proc Natl Acad Sci U.S.A 114:9641–9646. https://doi.org/10.1073/pnas.1707186114

15. Visôtto LE, Oliveira MGA, Guedes RNC, et al (2009) Contribution of gut bacteria to digestion and development of the velvetbean caterpillar, *Anticarsia gemmatalis*. J Insect Physiol 55:185–191. https://doi.org/10.1016/j.jinsphys.2008.10.017

16. Liang X, He J, Zhang N, et al (2022) Probiotic potentials of the silkworm gut symbiont *Enterococcus casseliflavus* ECB140, a promising L-tryptophan producer living inside the host. J Appl Microbiol 133:1620–1635. https://doi.org/10.1111/jam.15675

17. Shao Y, Chen B, Sun C, et al (2017) Symbiont-derived antimicrobials contribute to the control of the lepidopteran gut microbiota. Cell Chem Biol 24:66–75. https://doi.org/10.1016/j.chembiol.2016.11.015

18. Chen B, Mason CJ, Peiffer M, et al (2022) Enterococcal symbionts of caterpillars facilitate the utilization of a suboptimal diet. J Insect Physiol 138:104369. https://doi.org/10.1016/j.jinsphys.2022.104369

19. Mazumdar T, Hänniger S, Shukla SP, et al (2023) 8-HQA adjusts the number and diversity of bacteria in the gut microbiome of *Spodoptera littoralis*. Front Microbiol 14:1075557. https://doi.org/10.3389/fmicb.2023.1075557

20. Gadad H, Vastrad AS (2016) Gut bacteria mediated insecticide resistance in *Spodoptera litura*. J Exp Zool India 19:1099–1102

21. Gomes AFF, Omoto C, Cônsoli FL (2020) Gut bacteria of field-collected larvae of *Spodoptera frugiperda* undergo selection and are more diverse and active in metabolizing multiple insecticides than laboratory-selected resistant strains. J Pest Sci 93:833–851. https://doi.org/10.1007/s10340-020-01202-0

22. Li D, Zhang Y, Li W, et al (2019) Fitness and evolution of insecticide resistance associated with gut symbionts in metaflumizone-resistant *Plutella xylostella*. Crop Prot 124:104869. https://doi.org/10.1016/j.cropro.2019.104869

23. Li D-D, Li J-Y, Hu Z-Q, et al (2022) Fall Armyworm gut bacterial diversity associated with different developmental stages, environmental habitats, and diets. Insects 13:762. https://doi.org/10.3390/insects13090762

24. Shao Y, Arias-Cordero E, Guo H, et al (2014) In vivo Pyro-SIP assessing active gut microbiota of the cotton leafworm, *Spodoptera littoralis*. PLoS One 9(1):e85948. https://doi.org/10.1371/journal.pone.0085948

25. Tang X, Freitak D, Vogel H, et al (2012) Complexity and variability of gut commensal microbiota in polyphagous lepidopteran larvae. PLoS One 7(7):e36978. https://doi.org/10.1371/journal.pone.0036978

26. Xia X, Sun B, Gurr GM, et al (2018) Gut microbiota mediate insecticide resistance in the diamondback moth, *Plutella xylostella* (L.). Front Microbiol 9:25. https://doi.org/10.3389/fmicb.2018.00025

27. Vilanova C, Baixeras J, Latorre A, Porcar M (2016) The generalist inside the specialist: Gut bacterial communities of two insect species feeding on toxic plants are dominated by *Enterococcus* sp. Front Microbiol 7:1005. https://doi.org/10.3389/fmicb.2016.01005

28. Almeida LG, de Moraes LAB, Trigo JR, et al (2017) The gut microbiota of insecticide-resistant insects houses insecticide-degrading bacteria: A potential source for biotechnological exploitation. PLoS One 12(3):e0174754. https://doi.org/10.1371/journal.pone.0174754

29. Shao Y, Spiteller D, Tang X, et al (2011) Crystallization of α- and β-carotene in the foregut of *Spodoptera* larvae feeding on a toxic food plant. Insect Biochem Mol Biol 41:273–281. https://doi.org/10.1016/j.ibmb.2011.01.004

30. Higuita Palacio MF, Montoya OI, Saldamando CI, et al (2021) Dry and rainy seasons significantly alter the gut microbiome composition and reveal a key *Enterococcus* sp. (Lactobacillales: Enterococcaceae) core component in *Spodoptera frugiperda* (Lepidoptera: Noctuidae) corn strain from northwestern Colombia. J Pest Sci 21(6):10. https://doi.org/10.1093/jisesa/ieab076

31. Oliveira NC, Rodrigues PAP, Cônsoli FL (2022) Host-adapted strains of *Spodoptera frugiperda* hold and share a core microbial community across the western hemisphere. Microb Ecol. https://doi.org/10.1007/s00248-022-02008-6

32. Cappellozza S, Saviane A, Tettamanti G, et al (2011) Identification of *Enterococcus mundtii* as a pathogenic agent involved in the “flacherie” disease in *Bombyx mori* L. larvae reared on artificial diet. J Invertebr Pathol 106:386–393. https://doi.org/10.1016/j.jip.2010.12.007

33. Mason KL, Stepien TA, Blum JE, et al (2011) From commensal to pathogen: Translocation of *Enterococcus faecalis* from the midgut to the hemocoel of *Manduca sexta*. mBio 2(3):e00065–11. https://doi.org/10.1128/mBio.00065-11

34. Sun Y, Li X, Wang G, et al (2016) Genome sequence of *Enterococcus pernyi*, a pathogenic bacterium for the chinese oak silkworm, *Antheraea pernyi*. Genome Announc 4(3):e01764–15. https://doi.org/10.1128/genomeA.01764-15

35. Youngjin P, Kim K, Kim Y (2002) A pathogenic bacterium, *Enterococcus faecalis*, to the beet Armyworm, *Spodoptera exigua*. J Asia Pac Entomol 5:221–225. https://doi.org/10.1016/S1226-8615(08)60156-9

36. Devi S, Saini HS, Kaur S (2022) Assessing the pathogenicity of gut bacteria associated with tobacco caterpillar *Spodoptera litura* (Fab.). Sci Rep 12(1):8257. https://doi.org/10.1038/s41598-022-12319-w

37. Thakur A, Dhammi P, Saini HS, Kaur S (2015) Pathogenicity of bacteria isolated from gut of *Spodoptera litura* (Lepidoptera: Noctuidae) and fitness costs of insect associated with consumption of bacteria. J Invertebr Pathol 127:38–46. https://doi.org/10.1016/j.jip.2015.02.007

38. Rodrigues PAP, Oliveira NC, Omoto C, Girke T, Cônsoli FL (2023) Host plant affects the larval gut microbial communities of the generalist herbivores *Helicoverpa armigera* and *Spodoptera frugiperda*. BioRxiv 2023.03.08.531690; doi: https://doi.org/10.1101/2023.03.08.531690

39. Bauer AW, Kirby WM, Sherris JC, Turck M (1966) Antibiotic susceptibility testing by a standardized single disk method. Am J Clin Pathol 4:493–496. https://doi.org/10.1093/ajcp/45.4_ts.493

40. Andrews S. (2010) FastQC: a Quality Control Tool for High Throughput Sequence Data. In: Available online at: http://www.bioinformatics.babraham.ac.uk/projects/fastqc. http://www.bioinformatics.babraham.ac.uk/projects/fastqc

41. Bolger AM, Lohse M, Usadel B (2014) Trimmomatic: A flexible trimmer for Illumina sequence data. Bioinformatics 30:2114–2120. https://doi.org/10.1093/bioinformatics/btu170

42. Bankevich A, Nurk S, Antipov D, et al (2012) SPAdes: A new genome assembly algorithm and its applications to single-cell sequencing. J. Comput. Biol. 19:455–477. https://doi.org/10.1089/cmb.2012.0021

43. Prjibelski A, Antipov D, Meleshko D, et al (2020) Using SPAdes de novo assembler. Curr Protoc Bioinformatics 70(1):e102. https://doi.org/10.1002/cpbi.102

44. Brettin T, Davis JJ, Disz T, et al (2015) RASTtk: A modular and extensible implementation of the RAST algorithm for building custom annotation pipelines and annotating batches of genomes. Sci Rep 5(1):8365. https://doi.org/10.1038/srep08365

45. Alcock BP, Raphenya AR, Lau TTY, et al (2020) CARD 2020: Antibiotic resistome surveillance with the comprehensive antibiotic resistance database. Nucleic Acids Res 48:D517–D525. https://doi.org/10.1093/nar/gkz935

46. Wishart DS, Knox C, Guo AC, et al (2006) DrugBank: a comprehensive resource for in silico drug discovery and exploration. Nucleic Acids Res 34:D668–D672. https://doi.org/10.1093/nar/gkj067

47. Saier MH, Reddy VS, Moreno-Hagelsieb G, et al (2021) The transporter classification database (TCDB): 2021 update. Nucleic Acids Res 49:D461–D467. https://doi.org/10.1093/nar/gkaa1004

48. Chen L, Yang J, Yu J, et al (2005) VFDB: A reference database for bacterial virulence factors. Nucleic Acids Res 33:D325–D328. https://doi.org/10.1093/nar/gki008

49. Sayers S, Li L, Ong E, et al (2019) Victors: A web-based knowledge base of virulence factors in human and animal pathogens. Nucleic Acids Res 47:D693–D700. https://doi.org/10.1093/nar/gky999

50. Medema MH, Blin K, Cimermancic P, et al (2011) AntiSMASH: Rapid identification, annotation and analysis of secondary metabolite biosynthesis gene clusters in bacterial and fungal genome sequences. Nucleic Acids Res 39:W339–W346. https://doi.org/10.1093/nar/gkr466

51. Kautsar SA, Blin K, Shaw S, et al (2020) MIBiG 2.0: A repository for biosynthetic gene clusters of known function. Nucleic Acids Res 48:D454–D458. https://doi.org/10.1093/nar/gkz882

52. Meier-Kolthoff JP, Göker M (2019) TYGS is an automated high-throughput platform for state-of-the-art genome-based taxonomy. Nat Commun 10(1):2182. https://doi.org/10.1038/s41467-019-10210-3

53. Goris J, Konstantinidis KT, Klappenbach JA, et al (2007) DNA-DNA hybridization values and their relationship to whole-genome sequence similarities. Int J Syst Evol Microbiol 57:81–91. https://doi.org/10.1099/ijs.0.64483-0

54. Rosselló-Móra R, Amann R (2015) Past and future species definitions for Bacteria and Archaea. Syst Appl Microbiol 38:209–216. https://doi.org/10.1016/j.syapm.2015.02.001

55. Meier-Kolthoff JP, Hahnke RL, Petersen J, et al (2014) Complete genome sequence of DSM 30083T, the type strain (U5/41T) of *Escherichia coli*, and a proposal for delineating subspecies in microbial taxonomy. Stand Genomic Sci 9:2. https://doi.org/10.1186/1944-3277-9-2

56. Tamura K, Stecher G, Kumar S (2021) MEGA11: Molecular evolutionary genetics analysis version 11. Mol Biol Evol 38:3022–3027. https://doi.org/10.1093/molbev/msab120

57. Lefort V, Desper R, Gascuel O (2015) FastME 2.0: A comprehensive, accurate, and fast distance-based phylogeny inference program. Mol Biol Evol 32:2798–2800. https://doi.org/10.1093/molbev/msv150

58. Edgar RC (2004) MUSCLE: Multiple sequence alignment with high accuracy and high throughput. Nucleic Acids Res 32:1792–1797. https://doi.org/10.1093/nar/gkh340

59. Cock PJA, Antao T, Chang JT, et al (2009) Biopython: Freely available Python tools for computational molecular biology and bioinformatics. Bioinformatics 25:1422–1423. https://doi.org/10.1093/bioinformatics/btp163

60. Le SQ, Gascuel O (2008) An improved general amino acid replacement matrix. Mol Biol Evol 25:1307–1320. https://doi.org/10.1093/molbev/msn067

61. Sánchez-Romero MA, Casadesús J (2020) The bacterial epigenome. Nat Rev Microbiol 18:7–20. https://doi.org/10.1038/s41579-019-0286-2

62. Seong HJ, Han SW, Sul WJ (2021) Prokaryotic DNA methylation and its functional roles. J Microbiol 59:242–248. https://doi: 10.1007/s12275-021-0674-y

63. Stephens C, Reisenauer A, Wright R, et al (1996) A cell cycle-regulated bacterial DNA methyltransferase is essential for viability. Natl Acad Sci U S A. 93(3):1210–1214. https://doi: 10.1073/pnas.93.3.1210

64. Kahramanoglou C, Prieto AI, Khedkar S, et al (2012) Genomics of DNA cytosine methylation in *Escherichia coli* reveals its role in stationary phase transcription. Nat Commun 3(1):886. https://doi.org/10.1038/ncomms1878

65. Militello KT, Mandarano AH, Varechtchouk O, Simon RD (2014) Cytosine DNA methylation influences drug resistance in *Escherichia coli* through increased sugE expression. FEMS Microbiol Lett 350:100–106. https://doi: 10.1111/1574-6968.12299

66. Manso AS, Chai MH, Atack JM, et al (2014) A random six-phase switch regulates pneumococcal virulence via global epigenetic changes. Nat Commun 5(1):5055. https://doi.org/10.1038/ncomms6055

67. Li J, Li JW, Feng Z, et al (2016) Epigenetic switch driven by DNA inversions dictates phase variation in *Streptococcus pneumoniae*. PLoS Pathog 12(7):e1005762. https://doi.org/10.1371/journal.ppat.1005762

68. Oliver MB, Basu Roy A, Kumar R, et al (2017) *Streptococcus pneumoniae* TIGR4 phase-locked opacity variants differ in virulence phenotypes. mSphere 2(6):e00386–17. https://doi.org/10.1128/msphere.00386-17

69. Kumar S, Karmakar BC, Nagarajan D, et al (2018) N4-cytosine DNA methylation regulates transcription and pathogenesis in *Helicobacter pylori*. Nucleic Acids Res 46:3429–3445. https://doi.org/10.1093/NAR/GKY126

70. Estibariz I, Overmann A, Ailloud F, et al (2019) The core genome m5C methyltransferase JHP1050 (M.Hpy99III) plays an important role in orchestrating gene expression in *Helicobacter pylori*. Nucleic Acids Res 47:2336–2348. https://doi.org/10.1093/nar/gky1307

71. Mishra S, Lin Z, Pang S, et al (2021) Recent advanced technologies for the characterization of xenobiotic-degrading microorganisms and microbial communities. Front Bioeng Biotechnol 9:632095. https://doi.org/10.3389/fbioe.2021.632059

72. Russell RJ, Scott C, Jackson CJ, et al (2011) The evolution of new enzyme function: Lessons from xenobiotic metabolizing bacteria versus insecticide-resistant insects. Evol Appl 4:225–248. https://doi.org/10.1111/j.1752-4571.2010.00175.x

73. Bhatti S, Satyanarayana GNV, Patel DK, Satish A (2019) Bioaccumulation, biotransformation and toxic effect of fipronil in *Escherichia coli*. Chemosphere 231:207–215. https://doi.org/10.1016/j.chemosphere.2019.05.124

74. Gangola S, Bhatt P, Kumar AJ, et al (2022) Biotechnological tools to elucidate the mechanism of pesticide degradation in the environment. Chemosphere 296:133916. https://doi.org/10.1016/j.chemosphere.2022.133916

75. Sutherland TD, Horne I, Weir KM, et al (2004) Enzymatic bioremediation: From enzyme discovery to applications. Clin Exp Pharmacol Physiol 31(11):817–21. https://doi: 10.1111/j.1440-1681.2004.04088.x

76. Itoh H, Tago K, Hayatsu M, Kikuchi Y (2018) Detoxifying symbiosis: Microbe-mediated detoxification of phytotoxins and pesticides in insects. Nat Prod Rep 35(5):434–454. https://doi.org/10.1039/c7np00051k

77. Jaffar S, Ahmad S, Lu Y (2022) Contribution of insect gut microbiota and their associated enzymes in insect physiology and biodegradation of pesticides. Front Microbiol 13:979383. https://doi.org/10.3389/fmicb.2022.979383

78. Kikuchi Y, Hayatsu M, Hosokawa T, et al (2012) Symbiont-mediated insecticide resistance. Proc Natl Acad Sci U S A 109:8618–8622. https://doi.org/10.1073/pnas.1200231109

79. Cheng D, Guo Z, Riegler M, et al (2017) Gut symbiont enhances insecticide resistance in a significant pest, the oriental fruit fly *Bactrocera dorsalis* (Hendel). Microbiome 5:1–12. https://doi.org/10.1186/s40168-017-0236-z

80. Bhandari S, Poudel DK, Marahatha R, et al (2021) Microbial enzymes used in bioremediation. J Chem 2021:1–17. https://doi.org/10.1155/2021/8849512

81. Bose S, Kumar PS, Vo DVN (2021) A review on the microbial degradation of chlorpyrifos and its metabolite TCP. Chemosphere 283:131447. https://doi.org/10.1016/j.chemosphere.2021.131447

82. Sun R, Liu C, Zhang H, Wang Q (2015) Benzoylurea chitin synthesis inhibitors. J Agric Food Chem 63:6847–6865. https://doi.org/10.1021/acs.jafc.5b02460

83. Garrido-Sanz D, Manzano J, Martín M, et al (2018) Metagenomic analysis of a biphenyl-degrading soil bacterial consortium reveals the metabolic roles of specific populations. Front Microbiol 9:232. https://doi.org/10.3389/fmicb.2018.00232

84. Yu Y, Yin H, Peng H, et al (2020) Proteomic mechanism of decabromodiphenyl ether (BDE-209) biodegradation by *Microbacterium* Y2 and its potential in remediation of BDE-209 contaminated water-sediment system. J Hazard Mater 387:121708. https://doi.org/10.1016/j.jhazmat.2019.121708

85. Zhan H, Huang Y, Lin Z, et al (2020) New insights into the microbial degradation and catalytic mechanism of synthetic pyrethroids. Environ Res 182:109138. https://doi: 10.1016/j.envres.2020.109138

86. Zhu N, Li R, Zhang J, et al (2021) Photo-degradation behavior of seven benzoylurea pesticides with C3N4 nanofilm and its aquatic impacts on *Scendesmus obliquus*. Sci Total Environ 799:149470. https://doi.org/10.1016/j.scitotenv.2021.149470

87. Cleveland CB, Bormett GA, Saunders DG, et al (2002) Environmental fate of spinosad. 1. Dissipation and degradation in aqueous systems. J Agric Food Chem 50:3244–3256. https://doi.org/10.1021/jf011663i

88. Finkelstein ZI, Baskunov BP, Rietjens IMCM, et al (2001) Transformation of the insecticide teflubenzuron by microorganisms. J Environ Sci Health B 36:559–567. https://doi.org/10.1081/PFC-100106185

89. Lee Y, Jeong SE, Hur M, et al (2018) Construction and evaluation of a korean native microbial consortium for the bioremediation of diesel fuel-contaminated soil in Korea. Front Microbiol 9:2594. https://doi.org/10.3389/fmicb.2018.02594

90. Koga R, Moriyama M, Onodera-Tanifuji N, et al (2022) Single mutation makes *Escherichia coli* an insect mutualist. Nat Microbiol 7:1141–1150. https://doi.org/10.1038/s41564-022-01179-9

91. Simonsen AK (2022) Environmental stress leads to genome streamlining in a widely distributed species of soil bacteria. ISME Journal 16:423–434. https://doi.org/10.1038/s41396-021-01082-x

92. Antonelli P, Duval P, Luis P, et al (2022) Reciprocal interactions between anthropogenic stressors and insect microbiota. Environ Sci Pollut Res 29(43):64469–64488. https://doi.org/10.1007/s11356-022-21857-9

93. Hufnagel M, Koch S, Creti R, et al (2004) A putative sugar-binding transcriptional regulator in a novel gene locus in *Enterococcus faecalis* contributes to production of biofilm and prolonged bacteremia in mice. J Infect Dis 189(3):420–30. https://doi.org/10.1086/381150

94. Creti R, Koch S, Fabretti F, et al (2006) Enterococcal colonization of the gastro-intestinal tract: Role of biofilm and environmental oligosaccharides. BMC Microbiol 6(1):1–8. https://doi.org/10.1186/1471-2180-6-60

95. Klünemann M, Andrejev S, Blasche S, et al (2021) Bioaccumulation of therapeutic drugs by human gut bacteria. Nature 597:533–538. https://doi.org/10.1038/s41586-021-03891-8

96. Jindal S, Yang L, Day PJ, Kell DB (2019) Involvement of multiple influx and efflux transporters in the accumulation of cationic fluorescent dyes by *Escherichia coli*. BMC Microbiol 19(1):1–16. https://doi.org/10.1186/s12866-019-1561-0

97. Ahmed M, Lyass L, Markham PN, et al (1995) Two highly similar multidrug transporters of *Bacillus subtilis* whose expression is differentially regulated. J Bacteriol 177(14):3904–10. https://doi.org/10.1128/jb.177.14.3904-3910.1995

98. Gill MJ, Brenwald NP, Wise R (1999) Identification of an efflux pump gene, pmrA, associated with fluoroquinolone resistance in *Streptococcus pneumoniae*. Antimicrob Agents Chemother 43(1):187–9. https://doi.org/10.1128/aac.43.1.187

99. Schindler BD, Kaatz GW (2016) Multidrug efflux pumps of Gram-positive bacteria. Drug Resist Updat 27:1–13. https://doi.org/10.1016/j.drup.2016.04.003

100. Scharf ME, Wolfe ZM, Raje KR, et al (2022) Transcriptome responses to defined insecticide selection pressures in the German cockroach (*Blattella germanica* L.). Front Physiol 12:2570. https://doi.org/10.3389/fphys.2021.816675

101. Yu L, Yang H, Cheng F, et al (2021) Honey bee *Apis mellifera* larvae gut microbial and immune, detoxication responses towards flumethrin stress. Environ Pollut 290:118107. https://doi.org/10.1016/j.envpol.2021.118107

102. Dethlefsen L, McFall-Ngai M, Relman DA (2007) An ecological and evolutionary perspective on humang-microbe mutualism and disease. Nature 449:811–818. https://doi.org/10.1038/nature06245

103. Stoy KS, Gibson AK, Gerardo NM, Morran LT (2020) A need to consider the evolutionary genetics of host–symbiont mutualisms. J Evol Biol 33:1656–1668. https://doi.org/10.1111/jeb.13715

104. Mason CJ, Peiffer M, Chen B, et al (2022) Opposing growth responses of lepidopteran larvae to the establishment of gut microbiota. Microbiol Spectr 10(4):e01941–22. https://doi.org/10.1128/spectrum.01941-22

105. Verneuil N, Rincé A, Sanguinetti M, et al (2005) Contribution of a PerR-like regulator to the oxidative-stress response and virulence of *Enterococcus faecalis*. Microbiology 151(12):3997–4004. https://doi.org/10.1099/mic.0.28325-0

106. Michel A, Agerer F, Hauck CR, et al (2006) Global regulatory impact of ClpP protease of *Staphylococcus aureus* on regulons involved in virulence, oxidative stress response, autolysis, and DNA repair. J Bacteriol 188:5783–5796. https://doi.org/10.1128/JB.00074-06

107. de Diego-Diaz B, Treu L, Campanaro S, et al (2018) Genome sequence of *Enterococcus mundtii* EM01, isolated from *Bombyx mori* midgut and responsible for flacherie disease in silkworms reared on an artificial diet. Genome Announc 6(3):e01495–17. https://doi.org/10.1128/genomeA.01495-17

108. Audisio MC, Terzolo HR, Apella MC (2005) Bacteriocin from honey bee beebread *Enterococcus avium*, active against *Listeria monocytogenes*. Appl Environ Microbiol 71:3373– 3375. https://doi.org/10.1128/AEM.71.6.3373-3375.2005

109. Kawamoto S, Shima J, Sato R, et al (2002) Biochemical and genetic characterization of mundticin KS, an antilisterial peptide produced by *Enterococcus mundtii* NFRI 7393. Appl Environ Microbiol 68:3830–3840. https://doi.org/10.1128/AEM.68.8.3830-3840.2002

110. Liang X, Sun C, Chen B, et al (2018) Insect symbionts as valuable grist for the biotechnological mill: an alkaliphilic silkworm gut bacterium for efficient lactic acid production. Appl Microbiol Biotechnol 102:4951–4962. https://doi.org/10.1007/s00253-018-8953-1

111. Douglas AE (2017) The B vitamin nutrition of insects: the contributions of diet, microbiome and horizontally acquired genes. Curr Opin Insect Sci 23:65–69. https://doi.org/10.1016/j.cois.2017.07.012

112. Michalkova V, Benoit JB, Weiss BL, et al (2014) Vitamin B6 generated by obligate symbionts is critical for maintaining proline homeostasis and fecundity in tsetse flies. Appl Environ Microbiol 80:5844–5853. https://doi.org/10.1128/AEM.01150-14

113. Srinivasan R, Karaoz U, Volegova M, et al (2015) Use of 16S rRNA gene for identification of a broad range of clinically relevant bacterial pathogens. PLoS One 10(2):e117617. https://doi.org/10.1371/journal.pone.0117617

114. Fox, GE, Wisotzkey, JD, Jurtshuk, P (1992). How close is close: 16S rRNA sequence identity may not be sufficient to guarantee species identity. Int J Syst Bacteriol 166–170. https://doi.org/10.1099/00207713-42-1-166

115. Vandamme P, Pot B, Gillis M, et al (1996) Polyphasic Taxonomy, a Consensus Approach to Bacterial Systematics. Microbiol Rev 60(2):407–38. https://doi: 10.1128/mr.60.2.407-438

116. Vandamme P, Peeters C (2014) Time to revisit polyphasic taxonomy. Antonie van Leeuwenhoek 106:57–65. https://doi: 10.1007/s10482-014-0148-x

117. Chun J, Oren A, Ventosa A, et al (2018) Proposed minimal standards for the use of genome data for the taxonomy of prokaryotes. Int J Syst Evol Microbiol 68:461–466. https://doi.org/10.1099/ijsem.0.002516

